# A Photo-regulatable Intein Based Trans-Splicing tool for Protein and Organelle Relocalization to Different Subcellular Compartments

**DOI:** 10.1101/2025.07.20.665713

**Authors:** Cheng Tang, Qian Zhou, Ziqing Gao, Pei Peng, Bing Wang, Zuojun Liu, Shunqing Xu, Hanzeng Li

**Affiliations:** School of Environmental Science and Engineering, Hainan University, Haikou, China 570228

## Abstract

The dynamics of protein subcellular localization are intricately regulated, requiring new tools for controlling protein translocation to uncover the biological significance of post-translational regulation of protein localization. Here, we developed a new method, <u>P</u>rotein <u>R</u>erouting via <u>IN</u>tein-mediated <u>T</u>rans-<u>S</u>plicing (PRINTS), which enables precise control of protein translocation across diverse subcellular compartments. By reconstituting functional signaling peptides, PRINTS can efficiently relocalize fluorescent proteins to the 26S proteasome, nucleus, mitochondria, plasma membrane, endomembrane organelles, and liquid-liquid phase separation (LLPS) condensates. Furthermore, we incorporated the optically regulated dimerization domain CRY2clust into the PRINTS system, achieving light-mediated control of protein entry into the cell membrane and LLPS membraneless compartments. Strikingly, we observed that HNRNPA1 promiscuously recruit CRY2- or intein-containing proteins into LLPS condensate, whereas FXR1, another LLPS protein showed minimal non-specific activity. PRINTS also allow organelle relocalization, such as LLPS condensates or mitochondria to cell membrane in a light-controllable manner. Overall, PRINTS provides a versatile and robust platform for manipulating protein and organelle subcellular translocations, offering a powerful tool to investigate the regulatory coordination and crosstalk between membranous and membraneless compartments in response to physiological needs.

## Introduction

Eukaryotic cells are compartmentalized into membrane-bound and membraneless organelles that crosstalk and efficiently carry out specialized reactions(*1*). However, the mechanisms underlying metabolic exchange and crosstalk between these organelles remain largely unexplored(*2*). Protein translocations are tightly regulated as another layer of mechanism that fine-tunes protein activity post-translationally(*3*). A protein’s destination is often determined by its signal sequence, prior to full translation is completed. For example, proteins with an N-terminal secretion signal are typically translocated into the endoplasmic reticulum (ER) immediately after the signal peptide is synthesized(*4*). These signaling peptides, which direct subcellular localization, are often cleaved off once the protein reaches its target, preventing reversal of the process(*4*).

Organelle interactions are increasingly recognized as critical for coordinating metabolic fluxes and organelle dynamics(*5*, *6*). For example, mitochondrial DNA replication and subsequent fission often occur at ER-mitochondria contact sites(*7*). Coupling organelles is an emerging challenge in synthetic biology. Recent studies have shown that liquid-liquid phase separation (LLPS) condensate can sequester and release liquid droplet by tethering these two organelles using halo tags(*8*). This process restricts access of mitochondria to liquid droplet, thereby regulating cellular metabolism. However, no studies have yet been documented to bring other organelles into proximity for metabolic channeling or signaling transduction. While many cellular processes rely on the precise protein positioning, maneuvering protein translocation and even relatively small organelles across subcellular compartments remains a significant challenge for understanding their physiological roles(*9*).

Proteins containing inteins represent a rare group of protein that undergo protein splicing(*10*). After translation, the intein domain of a precursor protein possesses enzymatic activity that allows it to self-associate, cleave itself off, and ligate the flanking extein regions to form a covalent peptide bond(*11*, *12*). This process, known as cis-splicing, requires no external cofactors or energy input(*11*, *12*). Remarkably, when an intein is split into two parts, the two fragments can still reassemble and retain catalytic activity through a process referred to as trans-splicing(*13*). Studies have shown that asymmetric splitting of the intein can eliminate intrinsic affinity between fragments without compromising its enzymatic activity(*14*). Although intein-mediated protein-protein interaction is irreversible, in a way limiting its flexibility, it provides stronger hinge than other types of non-covalent links(*15*). This robustness may make it possible for manipulation of larger cargos, such as organelles for intracellular mobilization, overcoming the force of microtubules association. Intein has been harnessed to detect protein-protein interactions both *in vitro* and *in vivo*(*14*, *16*).

In this study, we developed a post-translational approach, termed Protein Rerouting via INtein-mediated Trans-Splicing (PRINTS), which enables controlled post-translational translocation of proteins to diverse subcellular compartments. PRINTS leverages split intein-mediated trans-splicing to reconstitute functional signaling peptides, directing proteins to the nucleus, mitochondria, 26S proteasome, plasma membrane, or membraneless organelles, such as liquid-liquid phase separation (LLPS) condensates (Fig. S1). We further enhanced PRINTS by incorporating optogenetic control through adding the CRY2clust dimerization domain, allowing light-induced regulation of protein translocation into plasma membrane or other organelles. Finally, we relocated mitochondria and LLPS condensates to plasma membrane via PRINTS.

## Results

### Protein degradation via intein-mediated degron reconstitution

We developed a method to target proteins for degradation by covalently reconstituting a degron through intein-mediated protein trans-splicing, directing the protein for poly-ubiquitination and subsequently sorted to 26S proteasome (Fig. 1A). To exemplify this design, we selected the hGemin degron, a well-characterized substrate of the anaphase promoting complex (APC), which undergoes reciprocal degradation with E3 ubiquitination ligase SCF^skp2^(*17*, *18*). As illustrated by Fig. 1B, hGeminin exhibit a cell cycle-dependent degradation pattern, being degraded in G1 phase and stabilized in S-G2-M phase. This oscillation property of hGeminin has been harnessed as one key component of the FUCCI sensor (hGeminin-hCdt1), which can monitor cell cycle progression *in situ*(*17*, *18*). To avoid constitutive degradation of degron tagged protein before trans-splicing, we split the hGeminin degron into two parts. The N-terminal half fused to the intein-N fragment, and the C-terminal half was fused to the intein-C fragment. The representative target protein, mCherry, was tagged with the hGeminin-N-intein-N fragment (Fig. 1A). This strategy ensures robust detection of protein instability changes. Before splicing, mCherry was stably expressed throughout the entire mitosis (Fig. 1B-E). Upon co-transfection of both fragments, the characteristic hGeminin degradation pattern was restored, indicating efficient degron reconstitution via trans-splicing, catalyzed by three tested inteins pairs (gp41-1, sspGyrB, and NrdJ1, Fig. 1B-E). We also devised a light inducible “ON-to-OFF” approach using a cell cycle-independent skp1 degron and Cry2clust, which forms a photo-inducible homodimer (Fig. 1F). Meanwhile, the gp41-1 intein was separated asymmetrically at the 105^th^ residue, which preserves the catalytic activity but eliminated the self-association property of intein(*14*). In this design, trans-splicing occurs only when the two fusion proteins were brought into proximity by Cry2clust dimerization upon blue light irradiation. As expected, attachment of the skp1 degron reduced target protein expression (Fig. 1G-H), though degradation is partial, likely due to strong overexpression in the transfection-based experimental setup. Conversely, in an “OFF-to-ON” approach (Fig. S2A), the intact skp1 degron did not induce constitutive degradation of the target protein, and its removal via photo-induced trans-splicing did not stabilize target protein expression as expected (Fig. S2B). This may result from continuous overexpression of transfected plasmids or the lower efficiency of the skp1 degron compared to hGeminin. These proof-of-concept experiments, using either the cyclic FUCCI sensor degron or the ubiquitous skp1 degron, demonstrated that split-intein mediated protein splicing is capable of modulating protein stability, offering a versatile tool for loss of function studies.

**Fig. 1.**
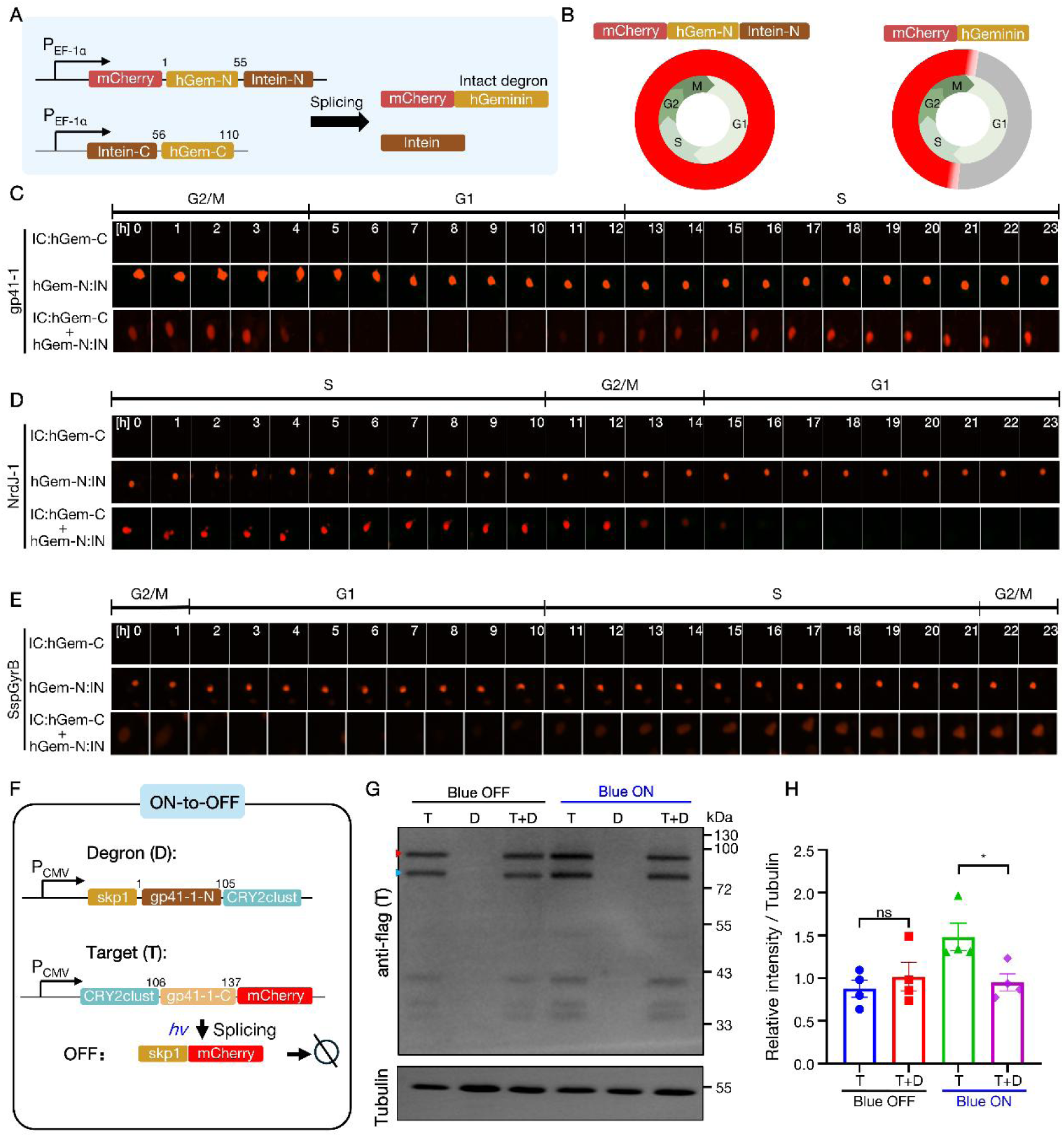
Intein-mediated reconstitution of degrons. **(A)** Schematic diagram of the plasmids encoding fusion proteins comprising split intein fragments and the human Geminin (hGeminin) degron that were derived from the FUCCI sensor, allowing cell cycle dependent protein degradation. The intein is split in a way that enables spontaneous ligation, while hGeminin degron is divided at the 55^th^ amino acid. **(B)** Anticipated fluorescence patterns during cell cycle, with indicated proteins aligned above the mitosis diagram. **(C-E)** Time-series images tracking a complete cell cycle in transfected cells expressing the proteins indicated on the left. Three intein pairs, *i.e.* gp41-1 (C), NrdJ-1 (D), and SspGyrB (E) were tested in independent experiments. A fully reconstituted, functional hGeminin degron confers periodic degradation during the G1 phase. **(F)** Schematic design of a light-controlled skp1 degron reconstitution. The gp41-1 intein was split at the 105^th^ amino acid, which preserved the splicing catalytic activity while eliminated self-association. Blue light (*hv*) irradiation triggers attachment of the skp1 derived degron to the target protein. **(G-H)** Evaluation of degradation efficiency for a representative target protein via Western blot. Quantification of relative protein levels, normalized to the loading control β-Tubulin is shown in H. The red triangle denotes band corresponding to the spliced protein, while the blue triangle indicates a non-specific band of unidentified splicing events. ns: not significant; *: *p*<0.05 but >0.01, as determined by *t*-test.

### Intein-mediated protein splicing enables nucleus or mitochondria translocation from cytosol

We next sought to determine if a similar strategy, *i.e.* reconstitution of split signaling peptides, could induce subcellular translocation of a given protein. To test this, we first split the SV40 nuclear localization signal (NLS) at the eighth amino acid, and fused the N-terminal and C-terminal sequences to split intein fragments respectively (Fig. 2A). The N-terminal parts of both intein and NLS were subsequently tagged to a fluorescent protein, mCherry. We found that the N-terminal of NLS *per se* did not promote nuclear localization of mCherry. However, co-expression with the C-terminal fragments resulted in robust nuclear localization, indicating efficient protein trans-splicing. This result demonstrated that post-translational reconstitution of signaling peptide via intein-mediated trans-splicing can effectively induce subcellular translocation. Although intein-mediated splicing leaves a six-amino acid scar (three from each extein) at the junction, these residues seem not to impair the functionality of the reconstituted NLS, as evidenced by its high efficiency.

**Fig. 2.**
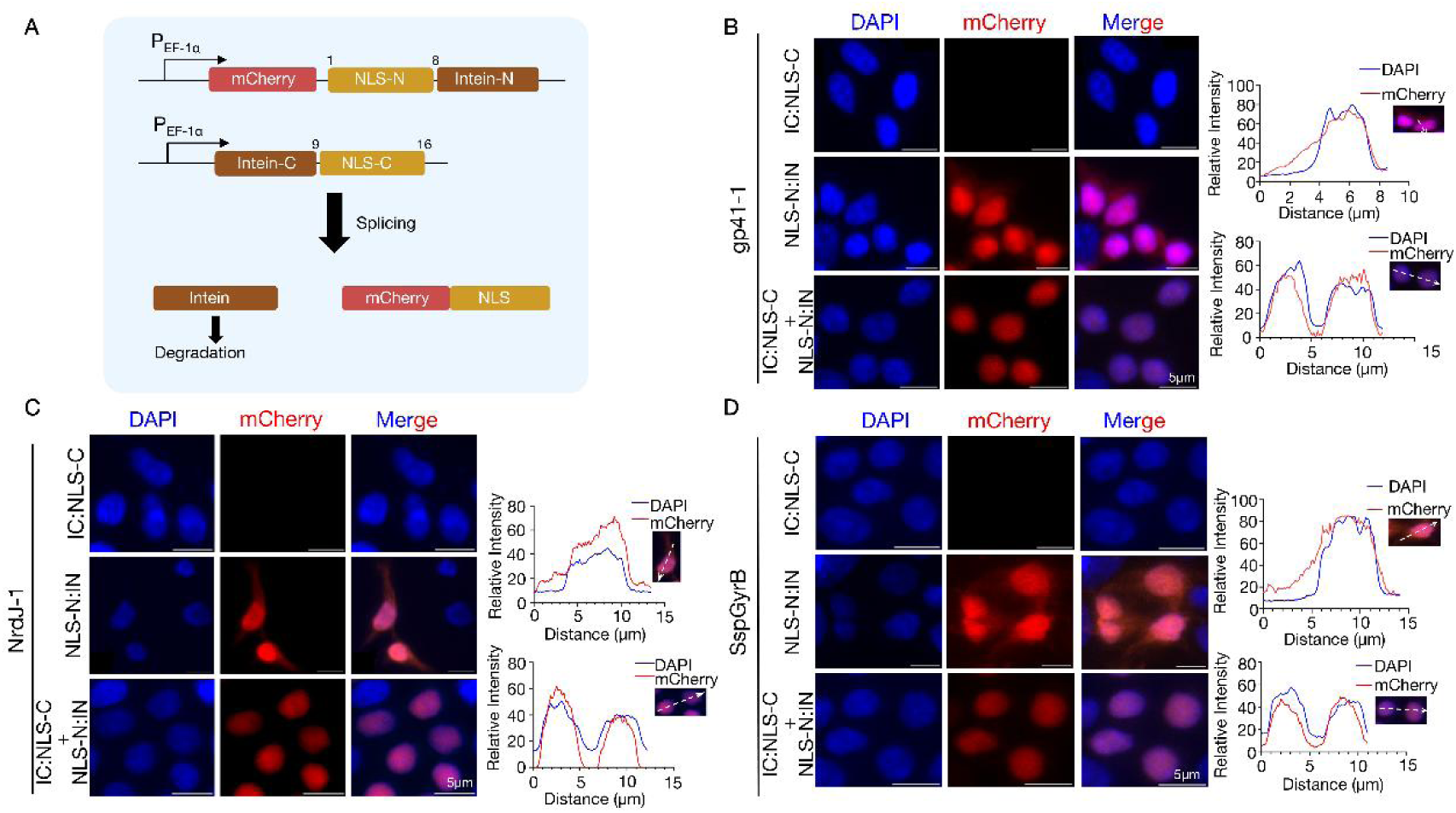
Controlled nucleus translocation via intein-mediated reconstitution of nuclear localization signal (NLS). **(A)** Schematic of the plasmid encoding proteins with split intein and NLS halves. The trans-splicing reaction generates a fusion protein with an intact NLS. The intein was divided into two fragments that allow spontaneous ligation. **(B-D)** Three intein pairs, *i.e.* gp41-1 (B), NrdJ-1 (C), and SspGyrB (D), were evaluated in proof-of-principle experiments. Representative images of HEK293T cells transfected with indicated plasmids were shown. Colocalization efficiency was analyzed by overlaying DAPI and mCherry fluorescence intensity profiles along manually drawn straight lines, as depicted to the right of each image. Scale bars represent 5 μm.

Next, we examined whether splicing a mitochondrial targeting signal (MTS) could also mediate post-translational translocation of a representative protein. Mitochondrial proteins are typically imported co-translationally through interaction with the TOMM complex, which translocate cytosolically synthesized proteins across the mitochondrial membranes, guided by the electrostatic properties of the MTS(*19*). In our initial design, we take use of the ATG4-derived MTS(*20*). It was tagged to the N-terminus of sfGFP, which is further fused with an intein-C parts at the N terminus of MTS (Fig. 3A&C). We hypothesized that the intein-C fragment would mask MTS activity, while co-expression with an intein-N fragment would trigger trans-splicing, removing the sequestering intein-C fragment to expose the MTS and direct the protein to mitochondria. This process would leave a 6-amino acid scar at the MTS N-terminus (Fig. 3A). We first evaluated whether these scar residues disrupt MTS function using intein tag from gp41-1, SspGyrB, and NrdJ-1 (Fig. 3A-B). Our results showed that none of the three scar sequences impaired ATG4 MTS mitochondrial targeting (Fig. 3B). However, the gp41-1 intein-C fragment failed to suppress MTS activity when fused at the N-terminus (Fig. 3C-D), suggesting that this design did not adequately block the MTS activity, unlike the endogenous ATG4 did(*20*). To remedy this, we took an alternative design and split the MTS into two halves, which resulted in a smaller tag to reduce the risk of mistargeting (Fig. 3E). As expected, the C-terminal MTS half, fused to any of the three intein-C fragments, insufficiently target sfGFP to mitochondria (Fig. 3F-H). Mitochondrial translocation occurred only in the presence of fusion protein consisting of the N-terminal MTS half and the intein-N fragment, *i.e.* MTS-N::intein-N (NrdJ-1 or SspGyrB). In contrast, gp41-1-mediated MTS reconstitution failed to recapitulate mitochondrial localization, suggesting that addition of the gp41-1 extein (SGYSSS) in the middle of MTS may interfere with MTS activity. These results suggest that NrdJ-1 and SspGyrB, but not gp41-1 inteins are compatible with split ATG4 MTS to restore mitochondrial targeting.

**Fig. 3.**
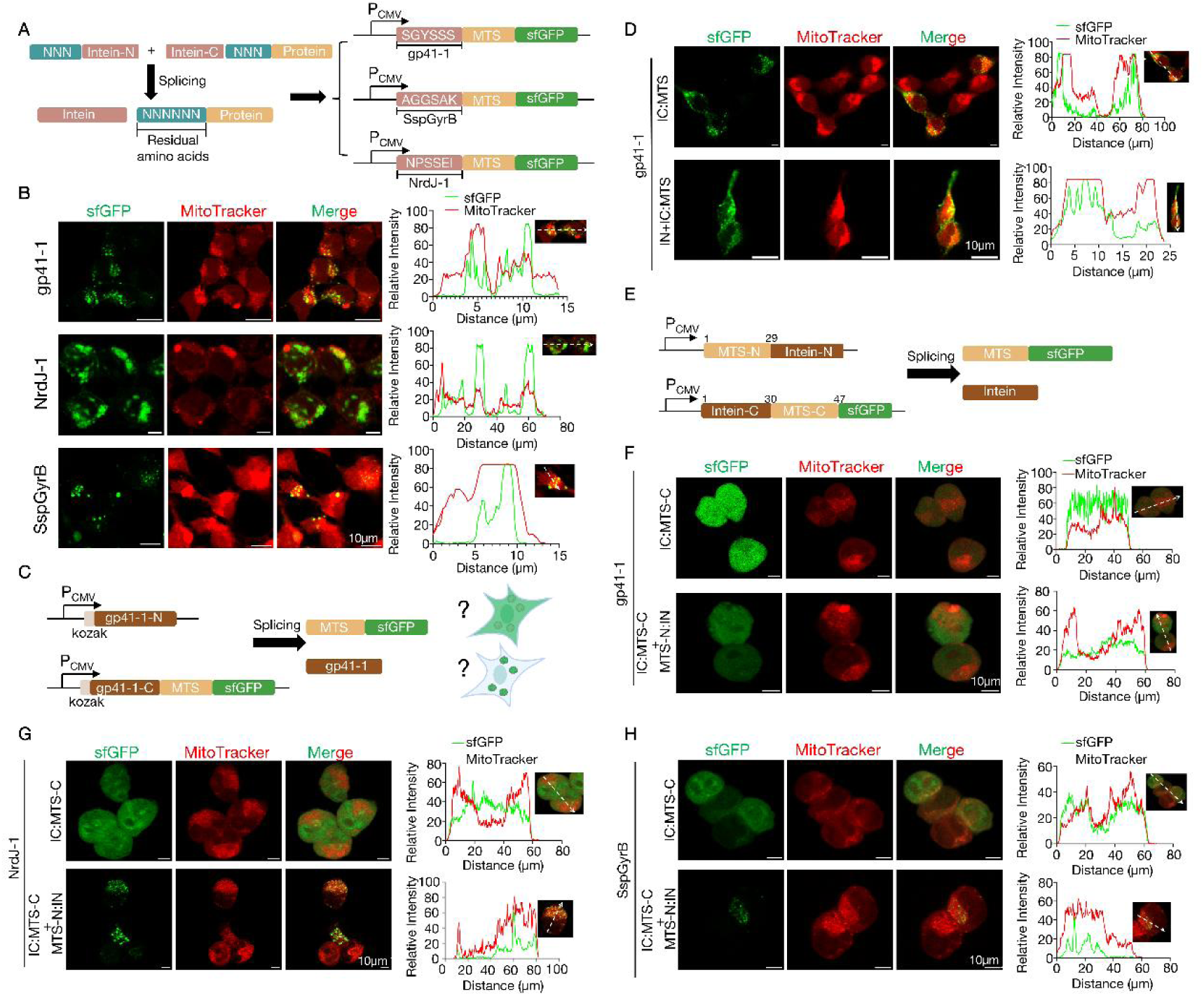
Protein translocation from cytoplasm to mitochondria via intein-mediated splicing. **(A)** Schematic of plasmids designed to assess the impact of residual extein amino acids post-splicing on mitochondrial targeting signal (MTS) efficacy. **(B)** Representative images of HEK293T cells transfected with the indicated plasmids (illustrated in A). Green fluorescence represents sfGFP, while red fluorescence marks mitochondria stained with MitoTracker Red. **(C)** Schematic diagram of plasmids that express split intein halves and split MTS. In this design, the intein C fragment was supposed to mask the mitochondria localization activity of MTS, while removal of the intein C domain by trans-splicing expose the MTS. The intein was split into two fragments in a way that allows spontaneous ligation. **(D)** Representative images of HEK293T cells transfected with the indicated plasmids (as illustrated in C). Scale bars represent 10µm. **(E)** Schematic of plasmids illustrating an alternative strategy that splits the MTS and reconstitutes a functional MTS by intein-mediated trans-splicing. **(F-H)** Representative images of cells transfected with the indicated plasmids. Three intein pairs with spontaneous capacity, *i.e.* gp41-1 (F), NrdJ-1(G), and SspGyrB (H) were trialed on this strategy. Colocalization analysis for images in B, D, F, G, and H were shown on the right. Scale bars represent indicated lengths in B, D, F, G, and H.

### Photo-inducible translocation of a cytosolic protein to plasma membrane or endomembrane system

Proteins localize to the plasma membrane either through transmembrane domains or post-translational prenylation(*21*, *22*). To achieve optogenetic control of protein translocation from the cytosol to the plasma membrane, we fused the target protein (sfGFP) to the N-terminal fragment of split gp41-1 (gp41-1-N, split at the 105^th^ amino acid) and the CRY2clust domain, which undergo dimerization upon blue light irradiation (Fig. 4A)(*23*). Additionally, we generated a chimeric anchor protein by fusing the CRY2clust to the C-terminal fragment of gp41-1 (gp41-1-C), mCherry and a plasma membrane localization signal (PMS) derived from Hras or Rac1(*24*). This latter fusion protein (anchor) displays sharp plasma membrane localization, with some expression in the Golgi apparatus. Before blue light induction, the target protein (sfGFP) was diffusely distributed throughout the cytosol. Upon blue light irradiation, it translocated to the plasma membrane or Golgi apparatus (Fig. 4B-C), likely initially driven by CRY2clust dimerization and subsequently stabilized through gp41-1 trans-splicing. When the PMS was shortened to a CAAX (where X represents any amino acid), the anchor protein predominantly localized to the endomembrane system (*e.g.* Golgi apparatus), consistent with prior reports (Fig. S3A)(*24*). As anticipated, the target protein shifted from the cytosol to endomembrane upon blue light irradiation (Fig. S3B-C). Thus, we demonstrated a cytosol-resident protein can be optogenetically directed to either the plasma membrane or endomembrane system via PRINTS.

**Fig. 4.**
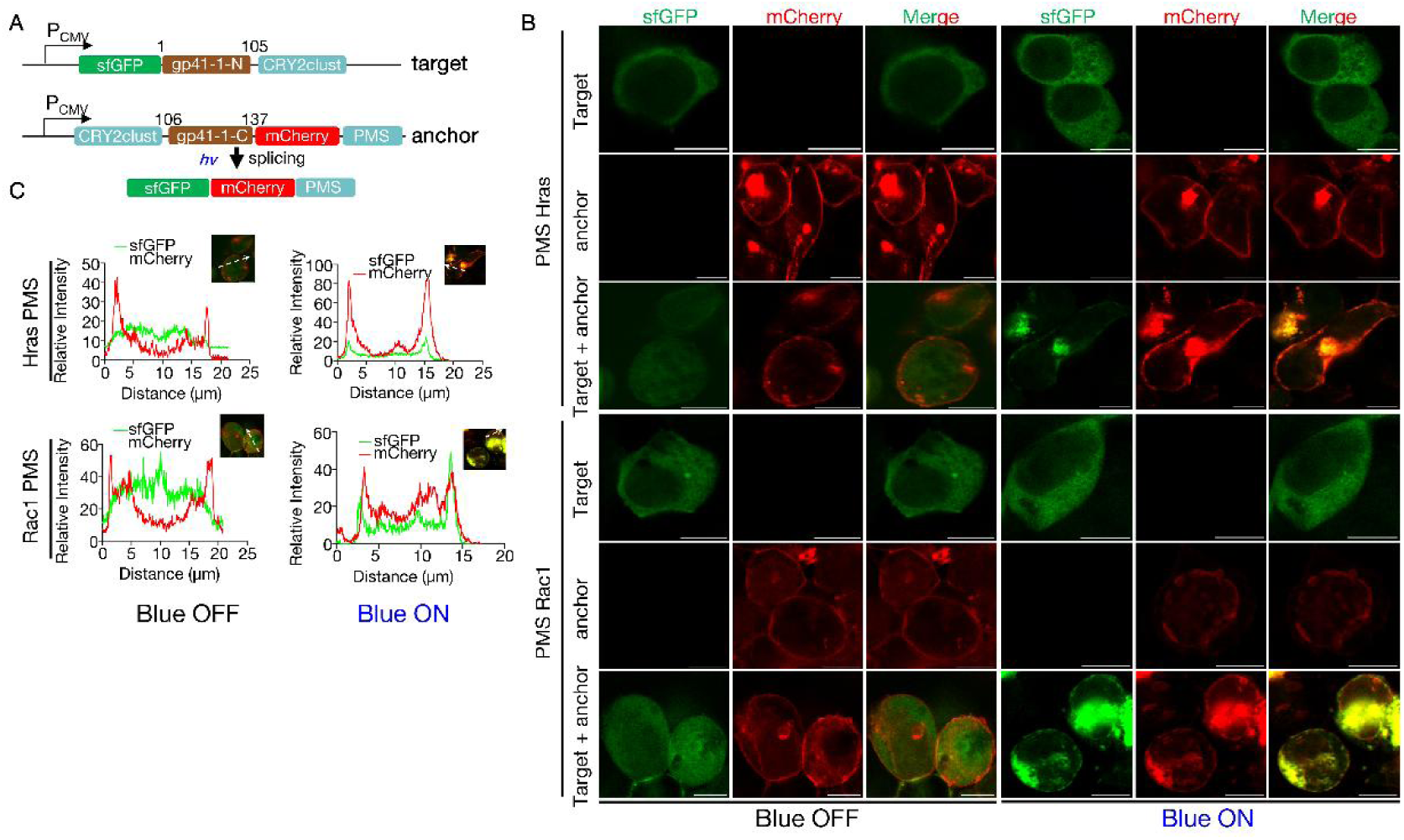
Optically controlled protein splicing enables inducible translocation of a cytosolic protein to the plasma membrane. **(A)** Schematic of plasmids designed to express the indicated proteins. The gp41-1 intein was split at the 105^th^ amino acid, which preserved the splicing catalytic activity while eliminated self-association. PMS denotes the <u>P</u>lasma <u>M</u>embrane Localization <u>S</u>ignal, comprising the CAAX sequence and a second membrane targeting signal upstream of CAAX, which were derived from Hras or Rac1 (see Table S1 for detailed amino acid sequences). *hv* represent blue light irradiation. **(B)** Representative images of HEK293T cells transfected with the plasmids shown in A. Cells were illuminated with 460 nm blue light for 30 minutes. Scale bars represent 10 μm. **(C)** colocalization efficiency analysis of images in B. Straight lines used for colocalization analysis are shown in the upper-right corner of each graph.

### Relocating a non-phase-separable protein to liquid-liquid phase separation subcellular condensates

Liquid-liquid phase separation (LLPS) organizes cellular processes by forming condensed microstructures comprising one or multiple types of proteins and nucleic acids(*25–27*). Key features of proteins capable of undergoing LLPS is the presence of intrinsically disordered regions (IDRs) or post-translational modifications, which facilitate multivalent protein-protein interaction or protein-nucleic acid interactions. Leveraging IDRs as LLPS recruitment signals, we applied auto-splicing split intein strategies to direct proteins to LLPS condensates. In our initial design, we used HNRNPA1, a protein known to spontaneously undergo LLPS, as an anchor site.

HNRNPA1 was fused to sfGFP at its N-terminus and to the N-terminal half of gp41-1 intein (split at the 93^rd^ amino acid, allowing for self-association and spontaneous splicing, Fig. 5A) at its C-terminus(*14*). Meanwhile, a representative target protein, mCherry, was tagged with the C-terminal gp41-1 intein and a FLAG tag for western blot detection of the splicing event (Fig. 5A). When expressed alone, the anchor protein (sfGFP::HNRNPA1) forms a LLPS condensate seemingly localized in the nucleus, while the target protein (gp41-1C::mCherry) were evenly diffused in the cytosol (Fig. 5B-C), which is consistent with expectation. Upon co-expression, the cytosolically expressed target protein was efficiently recruited to the HNRNPA1 LLPS condensate, as evidenced by colocalization of mCherry with sfGFP. Western blot analysis confirmed a robust trans-splicing event between anchor and target proteins (Fig. 5B).

**Fig. 5.**
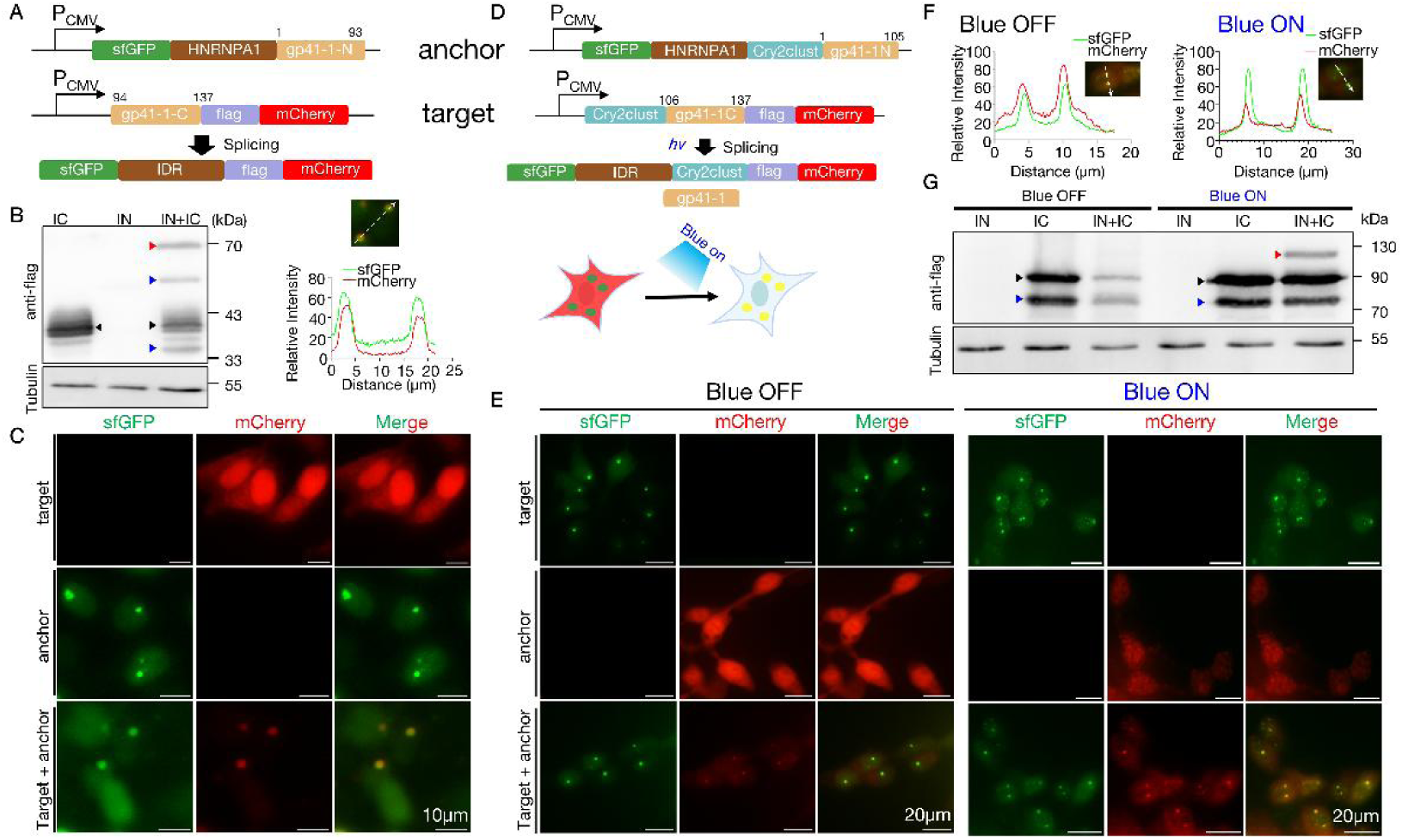
Relocalization of non-phase-separating proteins into HNRNPA1 liquid-liquid phase-separated subcellular condensates via protein splicing. **(A)** Schematic of the plasmids designed for localizing proteins to HNRNPA1 condensates using spontaneous splicing of gp41-1. The gp41-1 split after the E93 residue that enables spontaneous ligation. **(B)** Western blot validation of protein splicing events in HEK293T cells transfected with the indicated plasmids or their combinations. Samples loaded in each lane were labeled explicitly. The protein expressed by the IC plasmid and spliced product (illustrated in A) contains a FLAG tag used for Western blot analysis. **(C)** Representative images of sfGFP translocalization to HNRNPA1 condensates via spontaneous gp41-1 splicing. Colocalization analysis were shown above the images besides Fig. B. Scale bars represent 10 μm. **(D)** Schematic diagram of the plasmids for photo-regulatable relocation of non-phase-separating proteins to HNRNPA1 condensates. The gp41-1 intein was split at the 105^th^ amino acids, allowing splicing only when external forces bring the two halves into close proximity. Cry2clust, an optogenetic component that forms homodimer upon exposure to 460nm blue light. **(E-F)** Representative images of HEK293T cells transfected with the indicated plasmids, before and after illumination with 460nm blue light. Co-localization analysis of mCherry and the HNRNPA1 condensates (indicated by green fluorescence) were shown in F. Scale bars represent 20 μm. **(G)** Western blot validation of light-induced protein splicing events using anti-FLAG antibodies. Samples and treatment conditions were indicated above each lane. In b and g, black triangle denotes proteins expressed by the IC plasmid; red triangles indicate the recombination product after trans-splicing between IC and IN; blue triangles mark non-specific bands. β-Tubulin serves as the loading control.

To achieve optical control of protein translocation into the HNRNPA1 condensate, we took advantage of a previously mentioned asymmetrically split gp41-1 (Fig. 4A&5D). This version of split gp41-1 has minimal intrinsic affinity between its N-terminal and the C-terminal fragments, but retained splicing activity and only catalyze trans-splicing when brought into proximity(*14*). We incorporated the photo-regulatable CRY2clust domain, which homodimerizes via conformational alteration in response to blue light (460nm)(*23*). Cells expressing only the target protein (Cry2clust::gp41-1C::flag::mCherry) showed evenly distributed mCherry in the cytosol (Fig. 5E). Blue light exposure induced flake-like oligomeric structures of mCherry that is distinct from the HNRNPA1 LLPS condensates. In cells that express the chimeric anchor protein (sfGFP::HNRNPA1::cry2clust::gp41-1N fusion protein) formed a constitutive HNRNPA1 driven LLPS condensates (mostly 1-2 condensate *per* cell). Unexpectedly, present of both anchor and target protein in co-expression experiment resulted in high incidence of mCherry colocalization with sfGFP condensate regardless of blue light illumination (Fig. 5E-F). This suggests that either cry2clust or the C-terminal domain of gp41-1 has nonspecific binding affinity to HNRNPA1 LLPS condensates. However, the western blots results showed that trans-splicing occurs only upon blue light exposure, while the same transgenic cells in dark showed no trans-splicing between gp41-1 N- and C-terminal fragments despite they colocalized with each other (Fig. 5G). This indicates that cry2clust mediated protein dimerization is indispensable for trans-splicing, while co-existence of two intein halves in LLPS is not sufficient to drive trans-splicing reaction. To identify the source of non-specific binding, we tested two target constructs lacking either Cry2clust (Fig. S4A) or gp41-1C (Fig. S5A) from the original design (target protein of Fig. 5D). Both variants were evenly distributed in the cytosol in the dark, and formed blue light-induced oligomers (flake-like structures, and not a typical LLPS as distinguished by morphology) upon blue light irradiation as anticipated, when expressed alone. However, co-expression of the anchor protein led to constitutive recruitment of target proteins to HNRNPA1 LLPS condensates, regardless of blue light exposure (Fig. S4B-C and Fig. S5B-C). This indicates that both cry2clust and gp41-1C have intrinsic affinities to interact with HNRNPA1. The PONDR VSL2 disorder prediction analysis (https://www.pondr.com/) revealed that gp41-1C, but not Cry2clust is intrinsically disordered (Fig. S6). To minimize nonspecific effects from ambient light, we covered samples with aluminum foil and fixed them immediately before imaging. However, a study suggests that fixation may introduce artifacts, potentially altering LLPS condensates’ property(*28*). To exclude fixation artifacts, we imaged unfixed samples and still observed spontaneous recruitment of mCherry to HNRNPA1 LLPS condensates in dark (Fig. S5B-C). To determine if this nonspecific interaction is unique to HNRNPA1, or common to LLPS condensates, we tested FXR1, another well-characterized phase-separating protein. In the anchor construct, HNRNPA1 were replaced with FXR1. First, the sfGFP::FXR1 fusion protein constitutively formed droplet-like structures in the cytosol, but did not spontaneously recruit the target protein (CRY2clust::gp41-1C::FLAG::mCherry, Fig. S7A-B). Upon blue light exposure, the target protein formed flake-like oligomerization structure but did not colocalize with sfGFP::FXR1, as FXR1 lacks the cry2clust domain required for dimerization (Fig. S7B). Thus, FXR1 seems to be ideal for our purpose, as it eliminated nonspecific interactions between IDRs and Cry2clust domain or intein fragments. Next, we fused Cry2clust and gp41-1N to FXR1, which resulted in a chimeric protein sfGFP::FXR1::cry2clust::gp41-1N (anchor) that constitutively formed LLPS condensates as expected (Fig. 6A). Target protein remained evenly distributed in the cytosol without blue light, but formed flask-like structures upon blue light exposure when expressed alone (Fig. 6B-D). Eventually, co-expression with the anchor protein leads to a significant portion of mCherry being recruited to the FXR1 LLPS condensate under blue light, albeit to a lesser extent than HNRNPA1 (Fig. 6B-D). Furthermore, constructs lacking either the gp41-1C fragment (Fig. S8A-C) or CRY2clust domain (Fig. S9A-C) showed no obvious colocalization with FXR1 condensates. Consistently, western blot analysis confirmed that recombination only occurred in cells co-expressing anchor and target proteins upon blue light illumination (Fig. 6B). Thus, we have exemplified that PRINTS enables optogenetic manipulation of protein translocation into specific LLPS condensates by coupling Cry2clust dimerization with the following gp41-1 trans-splicing reaction.

**Fig. 6.**
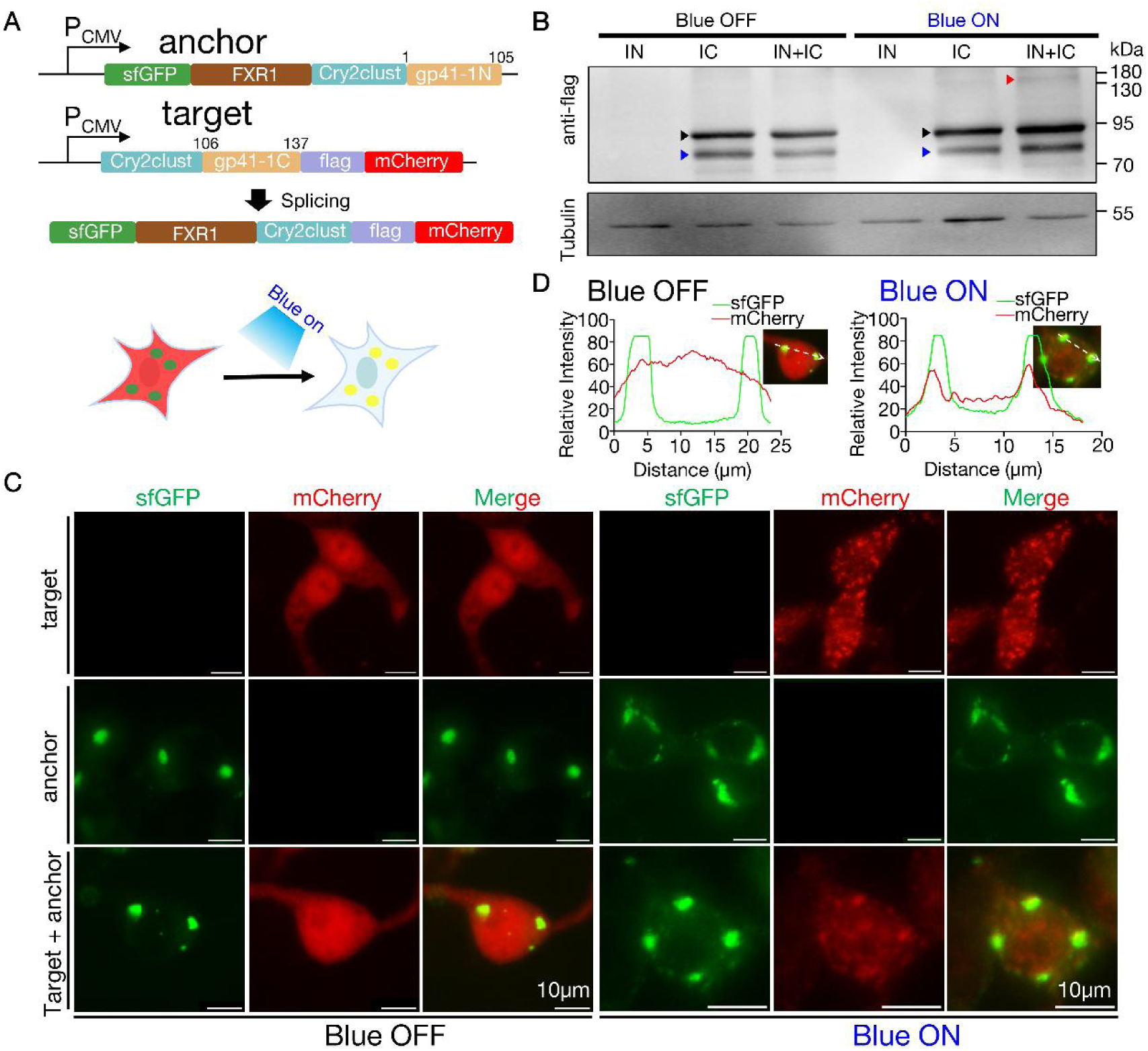
Photo-regulated translocation of a non-LLPS protein to FXR1-organized LLPS condensates. **(A)** Schematic of the plasmid used in the proof-of-concept experiment for light-controlled relocation to FXR1 LLPS condensates. **(B)** Western blot verification of light-induced protein trans-splicing events. Samples loaded in each lane were labeled explicitly. Black triangle denotes proteins expressed by the IC plasmid; red triangles indicate the recombination product after trans-splicing between IC and IN; blue triangles mark non-specific bands. β-Tubulin were used as the loading control. **(C-D)** Representative images of HEK293T cells transfected with the indicated plasmids, before and after irradiation with 460 nm blue light. Colocalization analysis of mCherry with FXR1 condensates (indicated by green fluorescence) were shown in D. Scale bars represent 10 μm.

### Photo-induced relocalization of LLPS condensates or mitochondria to plasma membranes or endomembranes

One key feature of the PRINTS tool distinct from other protein-protein interaction tools is its covalent linkage, rendering it irreversible. While this irreversibility may be a disadvantage compare to fast, reversible tools, it provides exceptional strength (covalent link), which makes it ideal for mobilizing large cargos such as organelles (Fig. S1). We thus interrogated whether PRINTS could tether two kinds of organelles that are otherwise distant, for example, LLPS condensates or mitochondria to the plasma membrane or Golgi apparatus. To test this, we co-expressed an LLPS targeting fusion protein (sfGFP::FXR1::Cry2clust::gp41-1N, termed “hook”, localized on the surface of LLPS condensate), and a plasma membrane-anchored fusion protein (Cry2clust::gp41-1C::mCherry::PMS, termed “anchor”, Fig. 7A). Before blue light-induced cry2clust dimerization, FXR1-constituted LLPS condensates were primarily cytosolic, distant from the plasma membrane or endomembranes like the Golgi apparatus (Fig. 7B-C). Upon blue light irradiation, FXR1 condensates become plasma membrane localized (Fig. 7B-C). The morphology of those membrane localized FXR1 signal are thick smears along the plasma membrane, likely due to fusion of individuals condensates during translocation. Similarly, HNRNPA1 condensates shifted from nuclear localization to a plasma membrane-associated pattern using the same experimental design (Fig. S10A-C). To confirm the LLPS nature of those plasma membrane-localized smears, we performed FRAP analysis, which revealed that these structures exhibit internal fluidity comparable to the endogenous FXR1 condensates (Fig. S11A-F). Substituting PMS with the shorter CAAX signal in the anchor protein showed similar plasma membrane translocation of LLPS condensates formed by HNRNPA1 (Fig. S12A-C) or FXR1 (Fig. S13A-C).

**Fig. 7.**
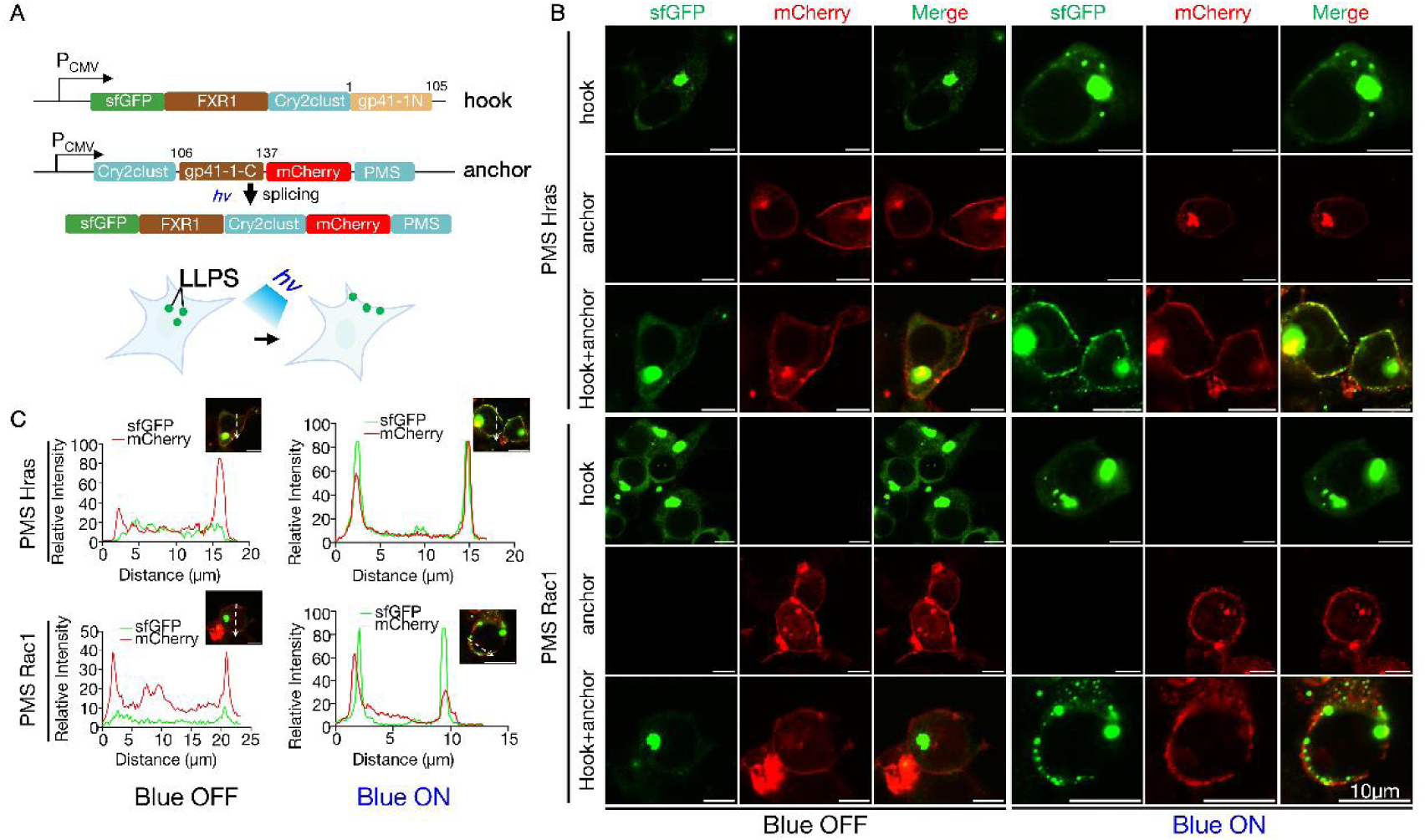
Photo-regulatable tethering of LLPS condensates (FXR1) to the plasma membrane. **(A)** Schematic of the plasmid used in a proof-of-concept experiment for light-controlled relocation of LLPS condensates (FXR1) to the plasma membrane via PRINTS mediated PMS covalent linking. **(B-C)** Representative images of HEK293T cells transfected with the indicated plasmids, captured before and after irradiation with 460 nm blue light (*hv*). Colocalization of green fluorescence (representing intact or dissembled LLPS condensates) with the plasma membrane and/or endomembrane system (marked by mCherry fluorescence) were shown in B. Quantitative colocalization analysis is presented in c. Scale bars represent 10 μm.

Next, we targeted mitochondria by anchoring a PRINTS component to the mitochondrial outer membrane via the Tomm20 signal peptide (tomm20::sfGFP::Cry2clust::gp41-1_N, hook), and nailing the second PRINTS component to the plasma membrane via PMS (Cry2clust::gp41-1_C::mCherry::PMS, anchor, Fig. 8A). Photo-inducible protein splicing relocated the sfGFP signal that represent mitochondria, to the peripheral region, colocalizing with the plasma membrane and endomembrane systems (Fig. 8B-C). A caveat of this observation is that only the mitochondria PRINTS fusion protein *per se* or a ruptured piece of mitochondria out membrane was brought to the plasma membrane. To confirm that the entire mitochondria, not just the PRINTS fusion protein or fragmented mitochondrial membranes were translocated, we co-stained cells with MitoTracker DeepRed FM, a mitochondrial matrix dye with distinct excitation and emission wavelengths from sfGFP and mCherry. Mitotracker DeepRed FM colocalize with the sfGFP signal at the plasma membrane, verifying whole-organelle translocation (Fig. 8D-E). Substituting PMS with the shorter CAAX signal in the anchor protein showed comparable results (Fig. S14A-C). Thus, PRINTS allows efficient, optically controlled tethering of LLPS condensates or mitochondria to the plasma membrane or endomembranes, demonstrating its potential for precise organelle relocalization.

**Fig. 8:**
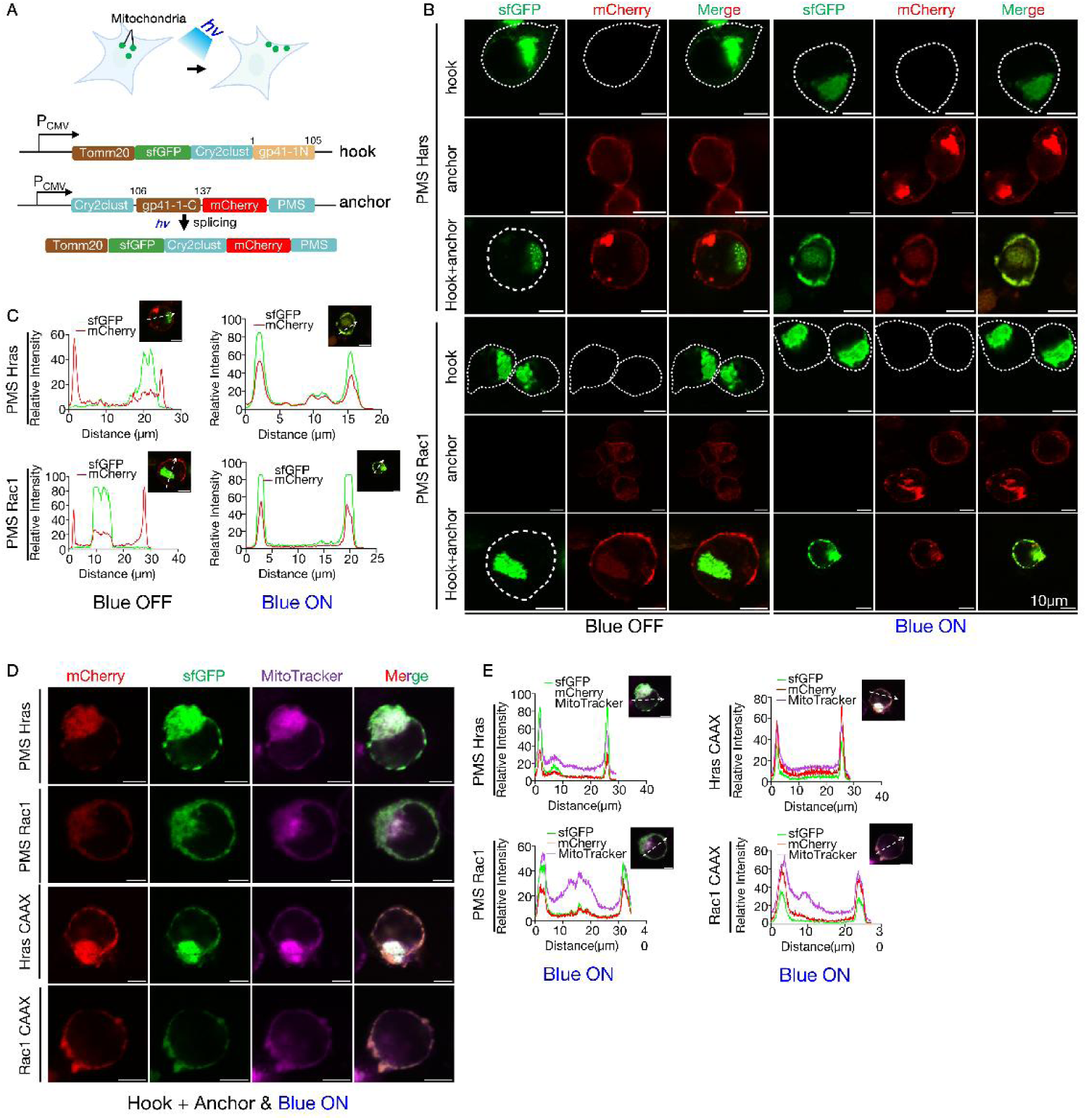
Light-controlled translocation of mitochondria to the plasma membrane using PRINTS. **(A)** Diagram of the plasmid used in the proof-of-concept experiment for optically controlled mitochondrial relocation to the plasma membrane via a plasma membrane localization signal (PMS). **(B-E)** Representative images of HEK293T cells transfected with the indicated plasmids, before and after irradiation with 460 nm blue light (*hv*). Colocalization of green fluorescence (mitochondria) with the plasma membrane and/or endomembrane system (marked by mCherry fluorescence) were shown. Colocalization analysis was shown in c. Mitochondria localized on plasma membrane after blue light irradiation were also stained with MitoTracker DeepRed FM and shown in D, while colocalization analysis were presented in E. Scale bars represent 10 μm.

## Discussion

In this study, we exemplified PRINTS through NLS, MTS, degron, PMS, and LLPS signal reconstitution, which provides a versatile and powerful tool for post-translationally relocalizing proteins within live cells. This offers a novel avenue to studying protein function and cellular organization. By reconstituting MTS from split fragments, we successfully directed proteins to these compartments with high efficiency. MTS often resides at the N-terminal of mitochondrial proteins, though exceptions exist for internal or C-terminal MTS. We unexpectedly found that embedding the ATG4 MTS in the middle of a fusion protein still result in mitochondrial targeting, suggesting that a special conformation of the ATG4 N-terminal fragment is required to sequester the ATG4 internal MTS (*20*).

LLPS has emerged as a fundamental mechanism in cellular organization, where proteins and RNA form membraneless organelles through multivalent interactions(*29–31*). These condensates play crucial roles in regulating multiple cellular events, however, their association with microtubes, or other organelles, and precise subcellular localization remain poorly understood due to limited tools for controlling protein entry into these structures(*32*, *33*). By coupling PRINTS with CRY2clust-based optogenetic control, we achieved light-responsive recruitment of non-phase separating proteins to LLPS condensates, which provides a spatial and temporal strategy to investigate their functional mechanisms and compositions when paired with the proximity labeling tools. In our first design, HNRNPA1 LLPS condensates nonspecifically include other proteins when CRY2clust was inactivated in the dark, even in the absence of binary binding moieties. In contrast, the FXR1 LLPS condensate exhibits better specificity. This raised important considerations regarding the specificity of LLPS condensate and protein interactions, emphasizing the need for careful selection of LLPS proteins for applications. Despite the constitutive colocalization of non-phase-separating in HNRNPA1 condensate, we observed robust and specific induction of protein trans-splicing only when CRY2clust dimerization is activated. This suggests that colocalization alone does not necessitate interaction, which is required for trans-splicing, implying the existence of a highly organized structure within the HNRNPA1 condensate. Although the physical chemistry of LLPS is not fully understood, evidences suggests that LLPS organizes its contents into layered structures(*34*, *35*). For example, in mitochondrial nucleoid assembly, the transcriptional factor A (TFAM) form an LLPS core with mtDNA, recruiting factors comprising the transcription initiation, elongation and termination complexes in a co-phase separation manner(*36*). These co-phase separation proteins form the shell of the LLPS droplet, surrounding the TFAM-mtDNA core and fulfilling both architectural and functional roles in the mitochondrial nucleoid LLPS droplet. He *et al* reported an assay for chromatin-bound condensates by exploratory sequencing (ACC-seq), which modulates condensate-mediated accessibility of DNA regions to the Tn5 transposase under native, fix, and 1,6-hex+fix conditions(*25*). In this assay, chimeric FUS(IDR)-SOX2 proteins form condensates at specific transcriptionally active DNA regions and are permeable to Tn5 transposase, whereas the addition of a fixative obstructs this accessibility. Phase-separating peptides were also recently explored as drug delivery coacervates for efficient macromolecules delivery(*37*).

These LLPS peptides are generally not selective in loading cargoes and can deliver proteins, DNA vectors, RNA, and small molecules to pass across the plasma membrane. Thus, the accessibility of LLPS condensates might depend on the composition of their core proteins, which can further partition within the LLPS condensates.

Intein-mediated trans-splicing forms covalent bonds, unlike reversible protein-protein interaction tools such as leucine zippers, halo-tags, or GFP-anti-GFP nanobodies(*11*, *38–40*). This covalent linkage limits its use in applications requiring reversible binding. However, it may enable the mobilization of larger cargos, such as organelles, which are often dynamically tethered to microtubules and need stronger force (longer OFF time) to prevent reversal transportation(*41*, *42*). Bring two organelles together can achieve multiple aims in synthetic biology, such as metabolic channeling(*43*). For example, tethering lipid droplet and mitochondria in adipose tissue could enhance the metabolism of lipids that may represent a prototype of curing obesity(*44*). Although not attempted in our study, tethering pathological aggregate that drive neurodegeneration to autophagosome-lysosome may promote its lysosomal degradation and slow down it the pathological processes.

In conclusion, PRINTS represents a useful tool in the field of protein manipulation, providing a highly customizable and efficient platform for the controlled translocation of proteins across subcellular compartments. The integration of optogenetic tools allows for real-time manipulation of protein/organelle localization, offering unprecedented flexibility for studying cellular organization, signal transduction, and phase separation.

## Acknowledgements

This study was supported by the National Science Foundation of China (32400724 and 22466015 to HL), Natural Science Foundation of Hainan Province (823QN230), Collaborative Innovation Center of One Health in Hainan University (XTCX2022JKA02 to SX), and Innovation Fund for Scientific and Technological Personnel of Hainan Province (KJRC2023B02 to SX).

## Author contributions

Conceptualization: H.L., and S.X. Methodology: C.T., H.L. Validation: H.L. Formal analysis: C.T. and H.L. Investigation: C.T., Q.Z., Z.G., P.P., B.W. Resources: H.L., and S.X. Writing—original draft: H.L. Writing—review and editing: H.L., and Z.L. Supervision: H.L. Project administration: H.L., and S.X. Funding acquisition: H.L., and S.X.

## Competing interests

The authors declare no conflict of interest.

## Methods

### Plasmids construction

All the plasmids made in this study and maps will be available upon request. The three pairs of inteins: gp41-1, NrdJ-1, and SspGyrB, HNRNPA1a IDR domain, and optogenetic element CRY2clust were chemically synthesized by Sango (Shanghai, China), according to published sequence information(*13*, *14*). We next cloned the indicated plasmids with each split intein fragment and localization signals into pcDNA3.1 (cut with NheI and EcoRI) or CSII-EF-mCherry-hGeminin backbone via Gibson Assembly method (NEBuilder-E2611, NEB, Boston). For hGeminin degrons, we PCRed split fragments of inteins (gp41-1, NrdJ-1, and SspGyrB), and divided hGeminin proteins (hGeminin_N, hGeminin_C) using primers with homologous overhangs, separately, then ligate intein half fragment and the corresponding hGeminin fragment into linearized CSII-EF-mCherry backbone that is a personal gift from Dr. Mingwei Min (Table S1). For NLS, we used a similar strategy and added two halves of NLSs (NLS-N or NLS-C) to the corresponding intein halves by PCR, and then ligated into the linearized CSII-EF-mCherry backbone using the Gibson Assembly method (Table S1).

To construct plasmids for proof-of-concept mitochondrial translocation, the coding sequences of the six extein residues (AGGSAK, NPCSEI, SGYSSS) for each intein was introduced as overhangs in the sense primers and PCRed the MTS fragment, which is subsequently ligated with sfGFP fragment and linearized pcDNA3.1 backbone (cut with NheI and EcoRI) via Gibson Assembly method (Table S1). The other gp41-1 intein halves were also PCRed with primers with overlap sequences and ligated with linearized pcDNA3.1 backbone and or sfGFP fragment via Gibson Assembly method (Table S1). After finding that gp41-1_C domain cannot mask the mitochondrial targeting effect of MTS, we decided to redesign the primers to divide MTS into two fragments with homologous arms (MTS-N and MTS-C). For this purpose, the indicated intein and MTS fragments were amplified by PCR using primers with homologous overlapping sequences and ligated with sfGFP coding fragment and linearized pcDNA3.1 backbone via Gibson Assembly method (Table S1).

For relocalizing a non-phase-separating protein into LLPS condensates, the HNRNPA1a IDR were PCRed from chemically synthesized DNA fragment, while the FXR1a genes were amplified from homemade human cDNA library. The intein halves that were separated as indicated in each figure, fluorescent proteins, IDR fragments, and/or cry2clust fragment were PCRed using primers with overlapping homologous sequences, and subsequently ligated by multiple fragment ligation into linearized pcDNA3.1 empty vector (cut with NheI and EcoRI) via the Gibson Assembly method (Table S1). Finally, to verify the domain requirement that mediates spontaneous entry into HNRNPA1a or FXR1 phase-separated droplets, we constructed plasmids lacking the cry2clust element and the gp41-1_C element by redesigning overlapping primers and PCRed corresponding fragments and ligated combination of fragments to linearized pcDNA3.1 empty vector via Gibson Assembly method (Table S1).

To make the target construct in the plasma membrane or endomembrane translocalization experiment, the sfGFP, gp41-1N half and the CRY2clust fragment were PCRed with primers with homologous overlapping sequences and ligated into linearized pcDNA3.1 vector (cut with NheI and EcoRI) via Gibson Assembly method. For the anchor plasmid, CAAX or plasma membrane localization signal were included as the 5’ overhanging sequences of anti-sense primer when amplifying the fluorescent protein fragment. The fluorescent proteins with C-terminal signal peptides, gp41-1-C, and CRY2clust fragments were PCRed and ligated to the same linearized pcDNA3.1 vector via Gibson Assembly method. All primers for cloning were listed in Table S2.

### Cell culture and transfection

HEK293T or Hela cells were cultured in DMEM (Gibco) medium supplemented with 10% fetal bovine serum (Newzerum) and 1% penicillin-streptomycin mixture, in a humidified incubator maintained at 37℃ and 5% CO2. The cells were tested using the Mycoplasma PCR Detection Kit (Beyotime) to confirm that there was no mycoplasma contamination. For imaging experiment, the cells were seeded onto 14-mm cell culture slides (Biosharp). Cells were grown to 40-60% confluence before being transfected with PEI (Sigma, 408727). The ratio between plasmid:PEI is 1:2.5. After a 48 hours culture in full medium post transfection, cells were imaged, or the protein were extracted for Western blot analysis.

### Western blots

In experiments where samples do not require blue-light irradiation, we collect proteins according to the normal protein-collection procedure. For the samples that need blue-light irradiation, we use repeated irradiations (irradiate for 5 minutes, then incubate for 1 hour, and repeat this cycle three times). The cold PBS washed cells were next suspended in RIPA lysis buffer (Solarbio) containing PMSF for lysis. After sonication on ice for 5 minutes, cell lysit were centrifugated at 15,000 rpm for 10 minutes at 4 °C. Subsequently, the supernatant was mixed with 4× protein loading buffer and incubated at 95 °C for 5 minutes. Proteins were separated by SDS-polyacrylamide gel electrophoresis (SDS-PAGE) and then transferred onto a polyvinylidene difluoride (PVDF) membrane (Millipore). The protein levels were determined by immunoblotting using an anti-Flag mouse monoclonal antibody (AF2852, Beyotime, China), and anti-β-Tubulin recombinant rabbit monoclonal antibody (ET1602-4, HUABIO, China).

### Time lapse imaging to measure the periodic degradation of hGeminin tagged fluorescent protein

The cells seeded in 12-well plates and grow to 50% confluence before being transfected with indicated plasmid(s). Fluorescence imaging was initiated 48 hours after transfection. The imaging was carried out using the Cytation One system (Agilent), with the imaging environment setting of 37°C and 5% CO₂. Images were captured every 30 minutes for a total imaging duration of 24 hours.

### Imaging of subcellular translocation of cytosolic fluorescent proteins

The cells were seeded in 6-well plates or a 14mm cell culture slides, and grow to an 50%-60% confluence before transfection. 48 hours post transfection, cells were stained with DAPI (Sigma), or Mitotracker Red or MitoTracker DeepRed FM (Thermo Fisher), and fixed with 4% paraformaldehyde for 5 minutes. Fluorescence images were acquired using the EVOS™ M7000 imaging system (Thermo Fisher), or a Nikon ECLIPSE Ci-L plus microscope.

### Blue light irradiation procedure

Cells were seeded onto 14mm cell culture slides, and grow to 80% confluence before transfection. 48hours after transfection, the cells in the non-blue-light illumination group were quickly fixed with 4% paraformaldehyde to avoid caveat caused by natural lights exposure. For the blue-light illumination group, 470-nm blue light was applied for 5 minutes before fluorescence imaging. For samples used for Western Blot analysis, the illumination time were appropriately extended to allow sufficient time for protein splicing.

### Colocalization coefficient quantification

All image analyses were conducted using ImageJ software (https://imagej.net/ij/). When performing colocalization analysis, the “Plot Profile” option within the “Analyze” menu of ImageJ was utilized to acquire the fluorescence intensity values at different positions along the drawn line. Subsequently, GraphPad Prism 8 was employed for graphing.

### Fluorescence recovery after photobleaching (FRAP)

The FRAP experiment was performed according to the method reported in Shi *et al* (*45*). FRAP were performed under a 60X oil-immersion objective in a Nikon AX confocal laser scanning microscope that is operated with NIS-Elements v6.0 software. A 488 nm laser at 10% power was used for bleaching, and fluorescence recovery was monitored every 3.5s for 180s post-bleach.

Fluorescence intensity in the bleached region was analyzed using ImageJ software.

### Statistical analysis

The presented data were obtained from a minimum of three independent experiments and were expressed as means ± standard deviation (SD) or standard error of mean (SEM). Statistical significance was determined with the student’s *t*-test, where *p*<0.05 and *p*<0.01 were represented as significant (*) and extremely significant (**) differences, respectively.

## Notes

### Competing Interest Statement

The authors have declared no competing interest.

## References

1. Y. G. Zhao, H. Zhang, Phase Separation in Membrane Biology: The Interplay between Membrane-Bound Organelles and Membraneless Condensates. Developmental Cell 55, 30–44 (2020).

2. Y. Liu, Y. Li, P. Zhang, Stress granules and organelles: coordinating cellular responses in health and disease. Protein Cell 16, 418–438 (2024).

3. E. Mills, K. Truong, Engineering Ca2+/calmodulin-mediated modulation of protein translocation by overlapping binding and signaling peptide sequences. Cell Calcium 47, 369–377 (2010).

4. Transfer of proteins across membranes. I. Presence of proteolytically processed and unprocessed nascent immunoglobulin light chains on membrane-bound ribosomes of murine myeloma. J Cell Biol 67, 835–851 (1975).

5. L. Zhou, M. Yu, M. Arshad, W. Wang, Y. Lu, J. Gong, Y. Gu, P. Li, L. Xu, Coordination Among Lipid Droplets, Peroxisomes, and Mitochondria Regulates Energy Expenditure Through the CIDE-ATGL-PPARα Pathway in Adipocytes. doi: 10.2337/db17-1452.

6. J.-F. Yang, X. Xing, L. Luo, X.-W. Zhou, J.-X. Feng, K.-B. Huang, H. Liu, S. Jin, Y.-N. Liu, S.-H. Zhang, Y.-H. Pan, B. Yu, J.-Y. Yang, Y.-L. Cao, Y. Cao, C. Y. Yang, Y. Wang, Y. Zhang, J. Li, X. Xia, T. Kang, R.-H. Xu, P. Lan, J.-H. Luo, H. Han, F. Bai, S. Gao, Mitochondria-ER contact mediated by MFN2-SERCA2 interaction supports CD8+ T cell metabolic fitness and function in tumors. Science Immunology 8, eabq2424 (2023).

7. S. C. Lewis, L. F. Uchiyama, J. Nunnari, ER-mitochondria contacts couple mtDNA synthesis with mitochondrial division in human cells. Science 353, aaf5549 (2016).

8. C. Amari, D. Simon, E. Pasquier, T. Bellon, M.-A. Plamont, S. Souquere, G. Pierron, J. Salvaing, A. R. Thiam, Z. Gueroui, Controlling lipid droplet dynamics via tether condensates. Nat Chem Biol, doi: 10.1038/s41589-025-01915-2 (2025).

9. C. S. C. Ng, A. Liu, B. Cui, S. M. Banik, Targeted protein relocalization via protein transport coupling. Nature 633, 941–951 (2024).

10. H. Wang, L. Wang, B. Zhong, Z. Dai, Protein Splicing of Inteins: A Powerful Tool in Synthetic Biology. Front Bioeng Biotechnol 10, 810180 (2022).

11. K. V. Mills, M. A. Johnson, F. B. Perler, Protein splicing: how inteins escape from precursor proteins. J Biol Chem 289, 14498–14505 (2014).

12. K. W. Y. Kwong, A. K. L. Ng, W. K. R. Wong, Engineering versatile protein expression systems mediated by inteins in Escherichia coli. Appl Microbiol Biotechnol 100, 255–262 (2016).

13. F. Pinto, E. L. Thornton, B. Wang, An expanded library of orthogonal split inteins enables modular multi-peptide assemblies. Nat Commun 11, 1529 (2020).

14. Z. Yao, F. Aboualizadeh, J. Kroll, I. Akula, J. Snider, A. Lyakisheva, P. Tang, M. Kotlyar, I. Jurisica, M. Boxem, I. Stagljar, Split Intein-Mediated Protein Ligation for detecting protein-protein interactions and their inhibition. Nat Commun 11, 2440 (2020).

15. H. M. Beyer, K. M. Mikula, M. Li, A. Wlodawer, H. Iwaï, The crystal structure of the naturally split gp41-1 intein guides the engineering of orthogonal split inteins from cis-splicing inteins. FEBS J 287, 1886–1898 (2020).

16. Z. Yao, J. Kim, B. Geng, J. Chen, V. Wong, A. Lyakisheva, J. Snider, M. R. Dimlić, S. Raić, I. Stagljar, A split intein and split luciferase-coupled system for detecting protein-protein interactions. Mol Syst Biol 21, 107–125 (2025).

17. G. D. Grant, K. M. Kedziora, J. C. Limas, J. G. Cook, J. E. Purvis, Accurate delineation of cell cycle phase transitions in living cells with PIP-FUCCI. Cell Cycle 17, 2496–2516 (2018).

18. L. Fischer, I. Thievessen, FUCCI Reporter Gene-Based Cell Cycle Analysis. Methods Mol Biol 2644, 371–385 (2023).

19. S. Callegari, N. S. Kirk, Z. Y. Gan, T. Dite, S. A. Cobbold, A. Leis, L. F. Dagley, A. Glukhova, D. Komander, Structure of human PINK1 at a mitochondrial TOM-VDAC array. Science 388, 303–310 (2025).

20. Z. Antón, G. Mullally, H. C. Ford, M. W. van der Kamp, M. D. Szczelkun, J. D. Lane, Mitochondrial import, health and mtDNA copy number variability seen when using type II and type V CRISPR effectors. J Cell Sci 133, jcs248468 (2020).

21. Q. Zeng, X. Si, H. Horstmann, Y. Xu, W. Hong, C. J. Pallen, Prenylation-dependent Association of Protein-tyrosine Phosphatases PRL-1, −2, and −3 with the Plasma Membrane and the Early Endosome*. Journal of Biological Chemistry 275, 21444–21452 (2000).

22. W. F. Wade, I. Khrebtukova, K. L. Schreiber, D. J. McKean, T. K. Wade, Truncated MHC class II cytoplasmic and transmembrane domains: Effect on plasma membrane expression. Molecular Immunology 32, 433–446 (1995).

23. H. Park, N. Y. Kim, S. Lee, N. Kim, J. Kim, W. D. Heo, Optogenetic protein clustering through fluorescent protein tagging and extension of CRY2. Nat Commun 8, 30 (2017).

24. E. Choy, V. K. Chiu, J. Silletti, M. Feoktistov, T. Morimoto, D. Michaelson, I. E. Ivanov, M. R. Philips, Endomembrane trafficking of ras: the CAAX motif targets proteins to the ER and Golgi. Cell 98, 69–80 (1999).

25. J. He, X. Huo, G. Pei, Z. Jia, Y. Yan, J. Yu, H. Qu, Y. Xie, J. Yuan, Y. Zheng, Y. Hu, M. Shi, K. You, T. Li, T. Ma, M. Q. Zhang, S. Ding, P. Li, Y. Li, Dual-role transcription factors stabilize intermediate expression levels. Cell 187, 2746–2766.e25 (2024).

26. S. Saurabh, T. N. Chong, C. Bayas, P. D. Dahlberg, H. N. Cartwright, W. E. Moerner, L. Shapiro, ATP-responsive biomolecular condensates tune bacterial kinase signaling. Sci Adv 8, eabm6570 (2022).

27. S. He, S. Wang, Y. Lin, Biomolecular phase separation research: Milestones, insights, and future trajectories. CSB 69, 4486–4499 (2024).

28. S. Irgen-Gioro, S. Yoshida, V. Walling, S. Chong, Fixation can change the appearance of phase separation in living cells. Elife 11, e79903 (2022).

29. L. Yang, Z. Zhang, P. Jiang, D. Kong, Z. Yu, D. Shi, Y. Han, E. Chen, W. Zheng, J. Sun, Y. Zhao, Y. Luo, J. Shi, H. Yao, H. Huang, P. Qian, Phase separation-competent FBL promotes early pre-rRNA processing and translation in acute myeloid leukaemia. Nat Cell Biol 26, 946–961 (2024).

30. J. Shorter, Phase separation of RNA-binding proteins in physiology and disease: An introduction to the JBC Reviews thematic series. J Biol Chem 294, 7113–7114 (2019).

31. H. Chen, Y. Cui, X. Han, W. Hu, M. Sun, Y. Zhang, P.-H. Wang, G. Song, W. Chen, J. Lou, Liquid-liquid phase separation by SARS-CoV-2 nucleocapsid protein and RNA. Cell Res 30, 1143–1145 (2020).

32. V. A. Volkov, A. Akhmanova, Phase separation on microtubules: from droplet formation to cellular function? Trends in Cell Biology 34, 18–30 (2024).

33. M. Sun, M. Jia, H. Ren, B. Yang, W. Chi, G. Xin, Q. Jiang, C. Zhang, NuMA regulates mitotic spindle assembly, structural dynamics and function via phase separation. Nat Commun 12, 7157 (2021).

34. S. Ye, A. P. Latham, Y. Tang, C.-H. Hsiung, J. Chen, F. Luo, Y. Liu, B. Zhang, X. Zhang, Micropolarity governs the structural organization of biomolecular condensates. Nat Chem Biol 20, 443–451 (2024).

35. U. Rana, K. Xu, A. Narayanan, M. T. Walls, A. Z. Panagiotopoulos, J. L. Avalos, C. P. Brangwynne, Asymmetric oligomerization state and sequence patterning can tune multiphase condensate miscibility. Nat Chem 16, 1073–1082 (2024).

36. Q. Long, Y. Zhou, H. Wu, S. Du, M. Hu, J. Qi, W. Li, J. Guo, Y. Wu, L. Yang, G. Xiang, L. Wang, S. Ye, J. Wen, H. Mao, J. Wang, H. Zhao, W.-Y. Chan, J. Liu, Y. Chen, P. Li, X. Liu, Phase separation drives the self-assembly of mitochondrial nucleoids for transcriptional modulation. Nat Struct Mol Biol 28, 900–908 (2021).

37. Y. Sun, S. Y. Lau, Z. W. Lim, S. C. Chang, F. Ghadessy, A. Partridge, A. Miserez, Phase-separating peptides for direct cytosolic delivery and redox-activated release of macromolecular therapeutics. Nat Chem 14, 274–283 (2022).

38. W. T. Pu, K. Struhl, The leucine zipper symmetrically positions the adjacent basic regions for specific DNA binding. Proc Natl Acad Sci U S A 88, 6901–6905 (1991).

39. J. R. Moll, S. B. Ruvinov, I. Pastan, C. Vinson, Designed heterodimerizing leucine zippers with a ranger of pIs and stabilities up to 10−15 M. Protein Sci 10, 649–655 (2001).

40. H. Wang, J. Liu, K. P. Yuet, A. J. Hill, P. W. Sternberg, Split cGAL, an intersectional strategy using a split intein for refined spatiotemporal transgene control in Caenorhabditis elegans. Proc Natl Acad Sci U S A 115, 3900–3905 (2018).

41. L. Benedetti, A. E. S. Barentine, M. Messa, H. Wheeler, J. Bewersdorf, P. De Camilli, Light-activated protein interaction with high spatial subcellular confinement. Proceedings of the National Academy of Sciences 115, E2238–E2245 (2018).

42. T. Mashita, T. Kowada, H. Yamamoto, S. Hamaguchi, T. Sato, T. Matsui, S. Mizukami, Quantitative control of subcellular protein localization with a photochromic dimerizer. Nat Chem Biol 20, 1461–1470 (2024).

43. E. M. Zhao, N. Suek, M. Z. Wilson, E. Dine, N. L. Pannucci, Z. Gitai, J. L. Avalos, J. E. Toettcher, Light-based control of metabolic flux through assembly of synthetic organelles. Nat Chem Biol 15, 589–597 (2019).

44. L. Cui, A. H. Mirza, S. Zhang, B. Liang, P. Liu, Lipid droplets and mitochondria are anchored during brown adipocyte differentiation. Protein Cell 10, 921–926 (2019).

45. Y. Shi, Y. Liao, Q. Liu, Z. Ni, Z. Zhang, M. Shi, P. Li, H. Li, Y. Rao, BRD4-targeting PROTAC as a unique tool to study biomolecular condensates. Cell Discov 9, 47 (2023).

