## Supplemental Information for "A Photo-regulatable Intein Based Trans-Splicing tool for Protein and Organelle Relocalization to Different Subcellular Compartments"

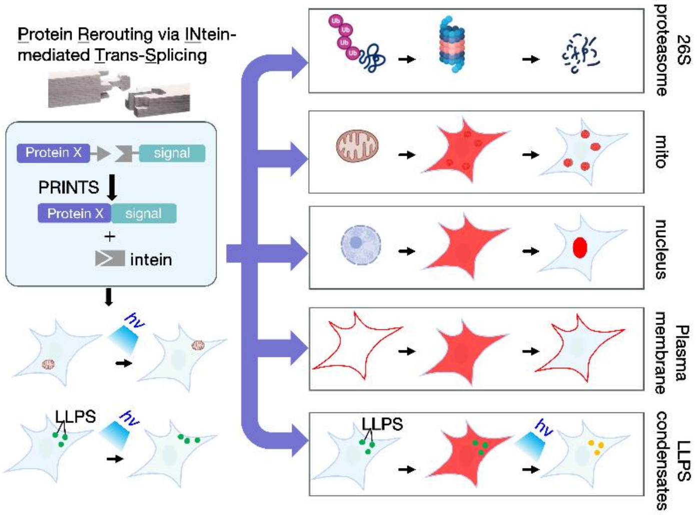


**Fig. S1. Schematic design of Protein Rerouting via INtein-mediated Trans-Splicing (PRINTS method) for relocalizing a protein or an organelle to various subcellular compartments using intein-mediated trans-splicing.** The diagram illustrates protein relocalization to subcellular locations, including the 26S proteasome, mitochondria (mito), nucleus, plasma membrane, and membraneless condensates organized via LLPS mechanism. *hv* represent photon.

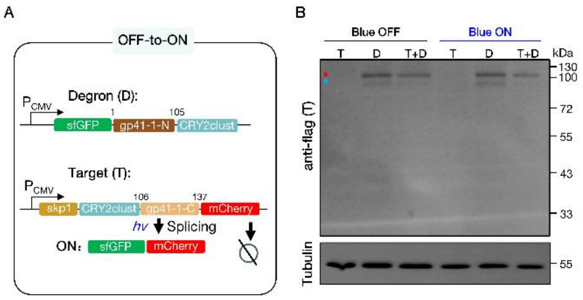


**Fig. S2. Protein trans-splicing did not achieve robust "OFF-to-ON" switch of a target protein. (A)** The "OFF-to-ON" design of a light-controlled removal of skp1 degron. The gp41-1 intein was split at the 105$^{th}$ amino acid, which preserved the splicing catalytic activity while eliminated self-association. Blue light irradiation triggers splicing that removes the skp1 degron from the target protein. **(B)** Evaluation of protein expression levels of a representative target protein via Western blot. The red triangle denotes band corresponding to the target protein, while the blue triangle indicates a non-specific band of unidentified splicing events.

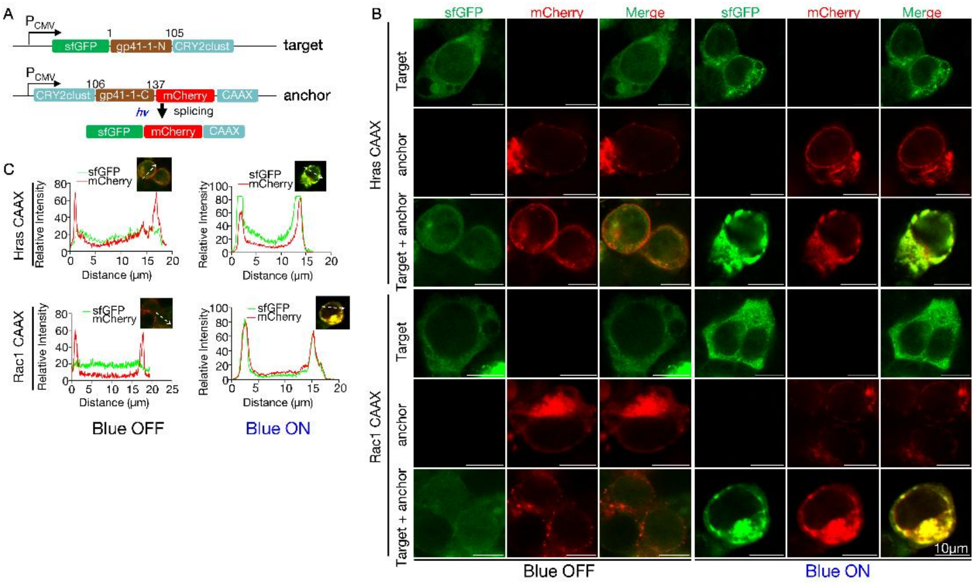


**Fig. S3. Optically controlled protein splicing allows inducible translocation of a cytosolic protein to endomembrane organelles. (A)** Schematic of plasmids designed to express the indicated proteins. The gp41-1 intein was split at the 105$^{th}$ amino acid, retaining the splicing catalytic activity while eliminating self-association. CAAX denotes the short signals for localization to endomembrane systems, such as Golgi apparatus. The CAAX were also derived from HRAS or Rac1 (see Table S1 for detailed amino acid sequences). *hv* represent blue light irradiation. **(B)** Representative images of HEK293T cells transfected with the plasmids shown in **A**. Cells were illuminated with 460 nm blue light for 30 minutes. Scale bars represent 10 μm. **(C)** Colocalization efficiency analysis of images in **B**. Straight lines used for colocalization analysis are shown in the upper-right corner of each graph.

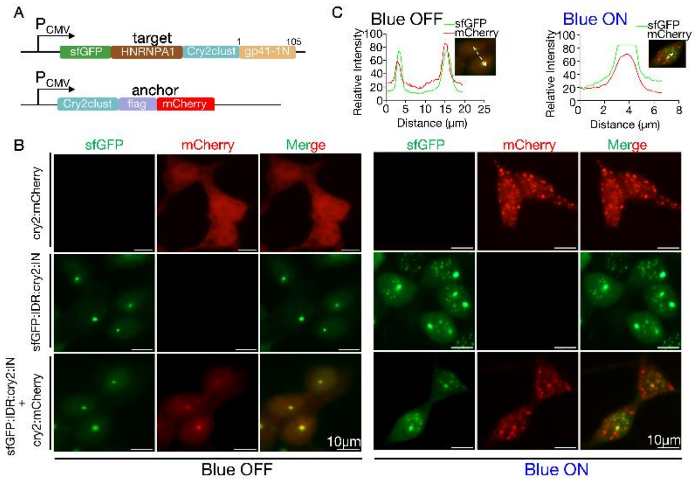


**Fig. S4. Optogenetics domain Cry2clust is constitutively recruited to HNRNPA1 LLPS condensates independent of light-induced conformational changes. (A)** Schematic of plasmids designed to determine the requirement of Cry2clust or gp41-1C for spontaneous entry into HNRNPA1 condensates. **(B-C)** Representative images of HEK293T cells transfected with the indicated plasmids, before (Blue OFF) and after blue light irradiation (460nm, labeled as Blue ON). Colocalization analysis between Cry2clust:flag::mCherry and HNRNPA1 condensates (green fluorescence) was shown in **C**. Scale bars represent 10 µm.

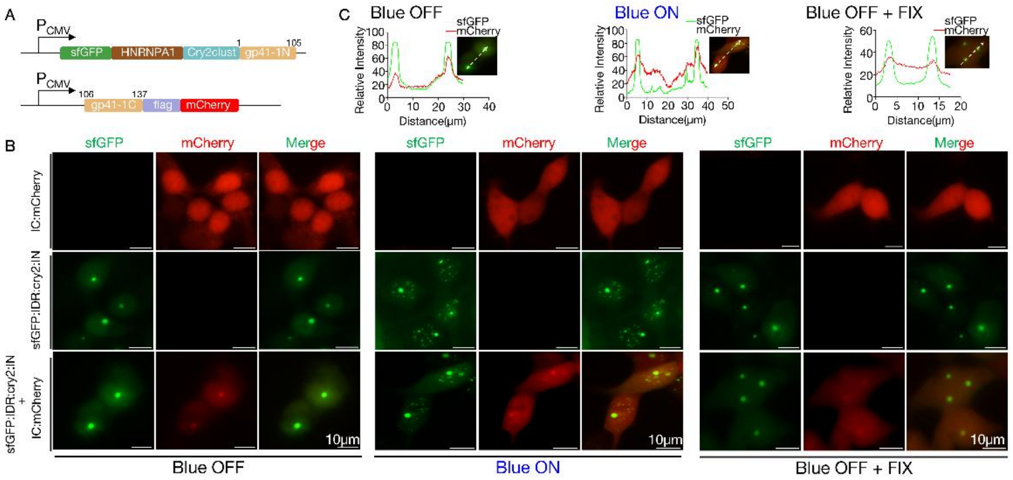


**Fig. S5. Spontaneous recruitment of the gp41-1 C-terminal domain into HNRNPA1 condensates. (A)** Schematic diagram of plasmids designed to verify whether Cry2clust or gp41-1C spontaneously mediate protein entry into HNRNPA1 condensates. **(B-C)** Representative images of HEK293T cells transfected with the indicated plasmids, before (Blue OFF), after blue light irradiation (Blue ON), or after fixation with 4% paraformaldehyde (Blue OFF+FIX). Colocalization analysis between Cry2clust:flag::mCherry and HNRNPA1 condensates (green fluorescence) was shown in **C**. Scale bars represent 10 µm.

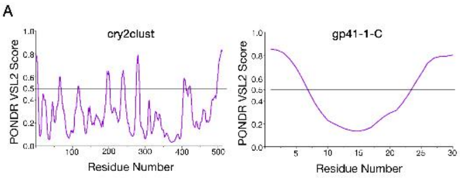


**Fig. S6. Prediction of intrinsically disordered regions in Cry2clust and gp41-1-C domains based on amino acid sequences. (A)** PONDR VSL2 scores for the indicated protein domains, predicted using the PONDR (Predictor of Natural Disordered Regions) online tool.

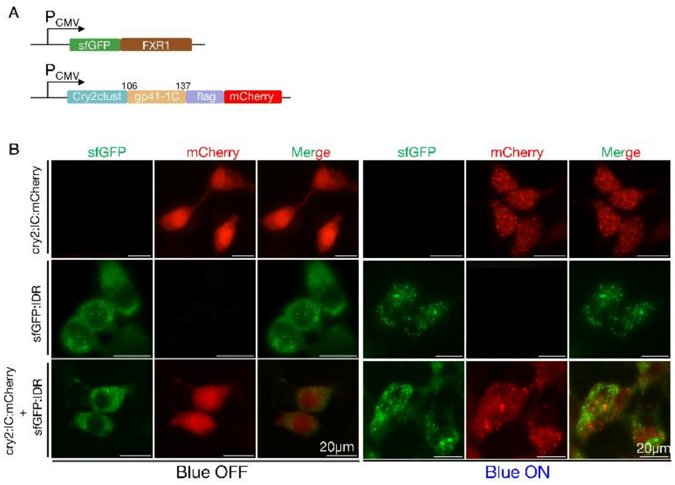


**Fig. S7. Neither the gp41-1C nor the cry2clust domain exhibit nonspecific recruitment to FXR1 LLPS condensates. (A)** Schematic diagram of plasmids designed to determine whether Cry2clust and gp41-1C spontaneously mediate protein entry into FXR1 condensates. **(B)** Representative images of HEK293T cells transfected with the indicated plasmids, before (Blue OFF) and after blue light irradiation (Blue ON).

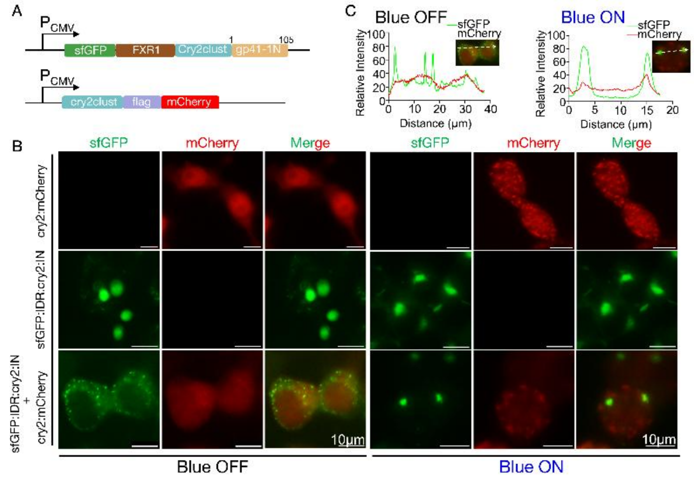


**Fig. S8. Photo-induced Cry2clust dimerization alone is not sufficient for protein translocation to FXR1 LLPS condensates. (A)** Schematic diagram of plasmids used to assess the efficiency of the optogenetic tag Cry2clust in promoting light-inducible entry of proteins into FXR1 condensates. **(B-C)** Representative images of HEK293T cells transfected with the indicated plasmids, before (Blue OFF) and after blue light irradiation (Blue ON). Colocalization analysis between Cry2clust:flag:mCherry fusion protein and FXR1 condensates (green fluorescence) was shown in **C**. Scale bars represent 10 μm.

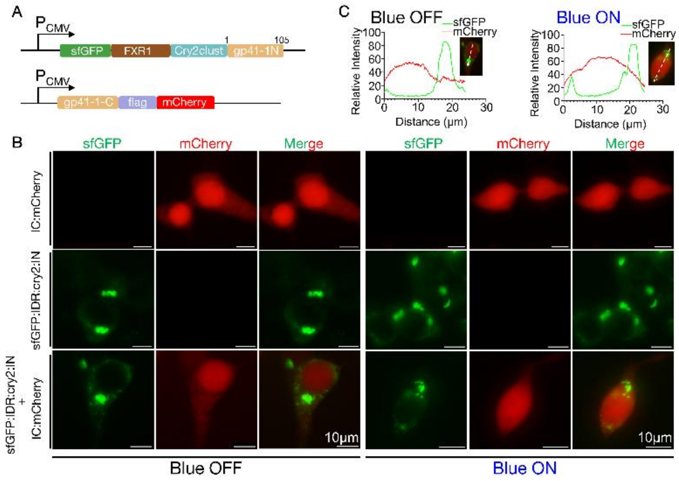


**Fig. S9. Non-spontaneous version of gp41-1 intein shows no non-specific recruitment into FXR1 LLPS condensates. (A)** Schematic of plasmids designed to assess whether gp41-1C constitutively mediates protein entry into FXR1 condensates. The gp41-1 intein was split after residue E105, which enables spontaneous ligation. **(B-C)** Representative images of HEK293T cells transfected with the indicated plasmids, before (Blue OFF) and after blue light irradiation (Blue ON). Colocalization analysis between gp41-1C:flag:mCherry fusion protein and FXR1 condensates (green fluorescence) was shown in **C**. Scale bars represent 10 µm.

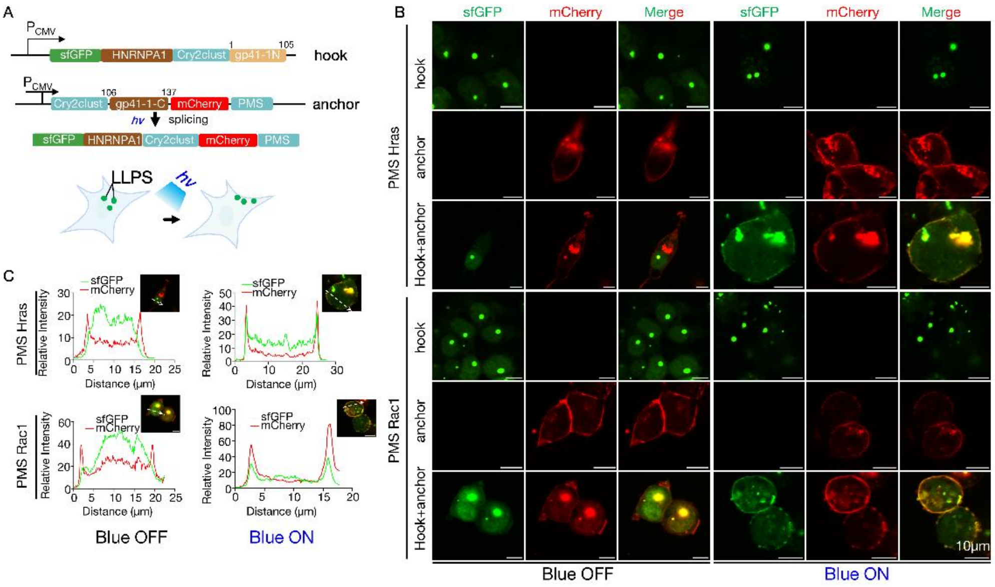


**Fig. S10. Photo-regulatable tethering of LLPS condensates (HNRNPA1) to the plasma membrane. (A)** Schematic of the plasmid used in a proof-of-concept experiment for light-controlled relocation of LLPS condensates (HNRNPA1) to the plasma membrane via PRINTS mediated PMS covalent linking. **(B-C)** Representative images of HEK293T cells transfected with the indicated plasmids, captured before and after irradiation with 460 nm blue light (*hv*). Colocalization of green fluorescence (representing intact or dissembled LLPS condensates) with the plasma membrane and/or endomembrane system (marked by mCherry fluorescence) were shown in B. Quantitative colocalization analysis is presented in C. Scale bars represent 10 μm.

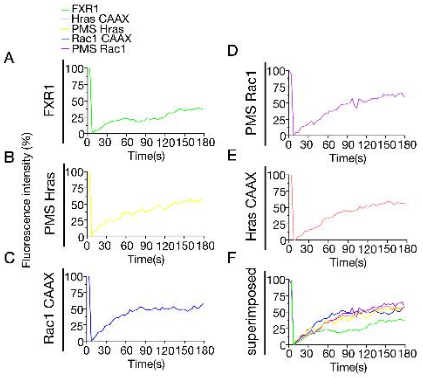

**Fig. S11. Fluorescence recovery after photobleaching (FRAP) analysis of LLPS condensates at varied locations. (A-F)** Fluorescence recovery curves of indicated LLPS condensates after photobleaching. FXR1 condensate (A), plasma membrane relocalized LLPS condensates (B-E) were overlayed to compare their internal fluidity and shown in F.

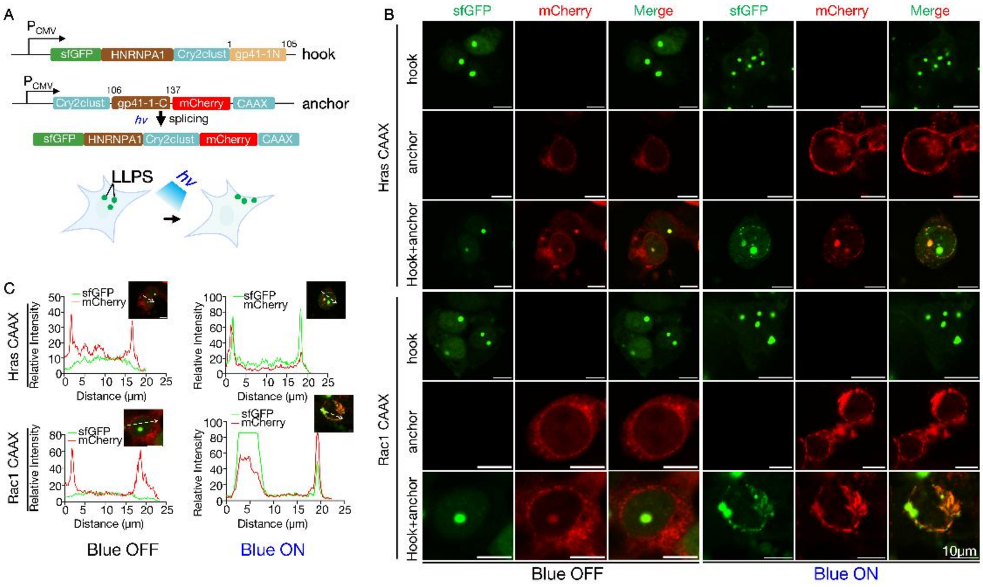

**Fig. S12. Photo-regulatable tethering of LLPS condensates (HNRNPA1) to the endomembrane system. (A)** Schematic of the plasmid used in a proof-of-concept experiment for light-controlled relocation of LLPS condensates (HNRNPA1) to the endomembrane system via PRINTS mediated CAAX covalent linking. **(B-C)** Representative images of HEK293T cells transfected with the indicated plasmids, captured before and after irradiation with 460 nm blue light (*hv*). Colocalization of green fluorescence (representing intact or dissembled LLPS condensates) with the endomembrane system (marked by mCherry fluorescence) were shown in B. Quantitative colocalization analysis is presented in C. Scale bars represent 10 µm.

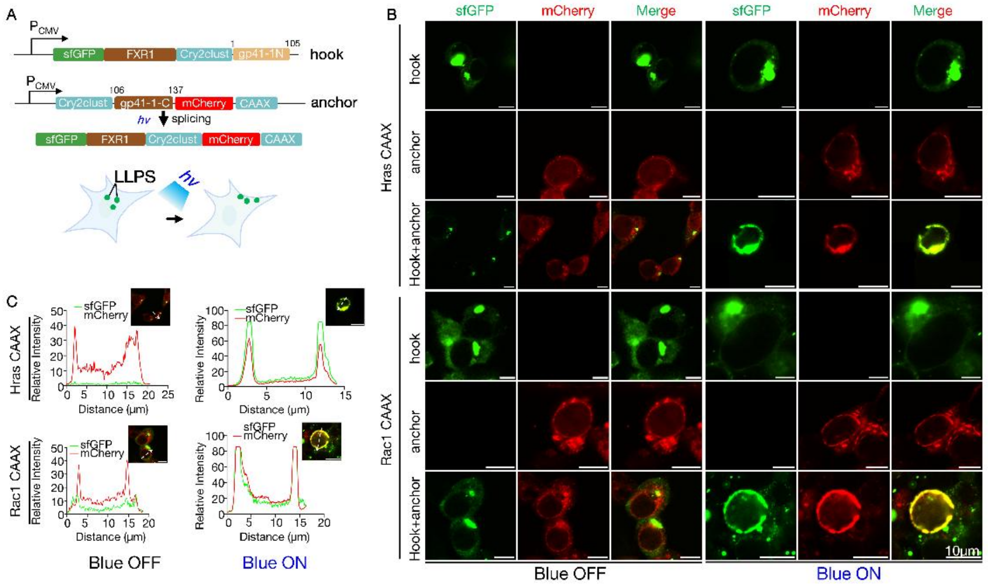


**Fig. S13. Photo-regulatable tethering of LLPS condensates (FXR1) to the endomembrane system. (A)** Schematic of the plasmid used in a proof-of-concept experiment for light-controlled relocation of LLPS condensates (HNRNPA1) to the endomembrane system via PRINTS-mediated CAAX covalent linking. **(B-C)** Representative images of HEK293T cells transfected with the indicated plasmids, captured before and after irradiation with 460 nm blue light (*hv*). Colocalization of green fluorescence (representing intact or dissembled LLPS condensates) with the endomembrane system (marked by mCherry fluorescence) were shown in B. Quantitative colocalization analysis is presented in C. Scale bars represent 10 µm.

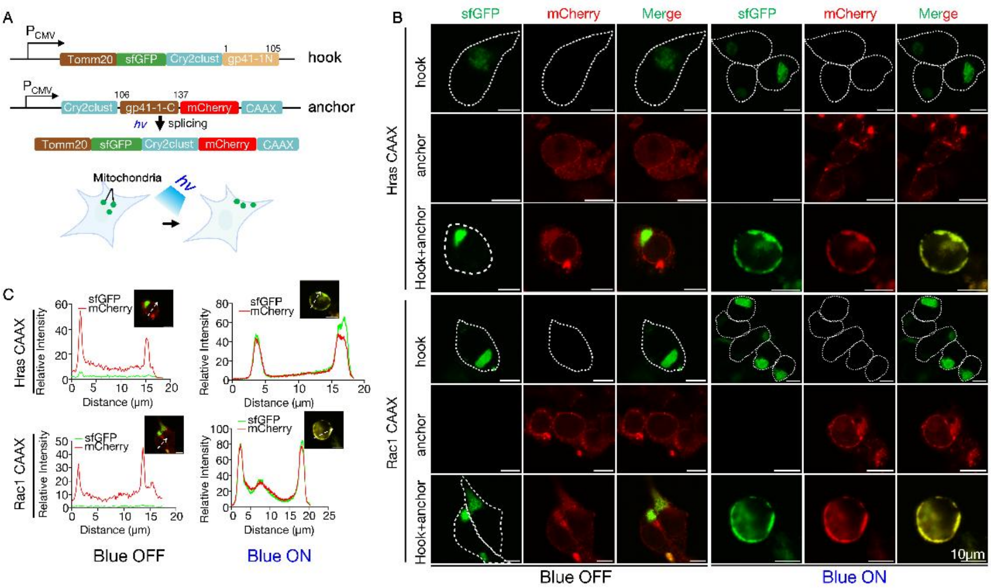


**Fig. S14. Light-controlled translocation of mitochondria to the endomembrane system using PRINTS. (A)** Diagram of the plasmid used in the proof-of-concept experiment for optically controlled mitochondrial relocation to the endomemrbanes via a CAAX signal. **(B-C)** Representative images of HEK293T cells transfected with the indicated plasmids, before and after irradiation with 460 nm blue light (*hv*). Colocalization of green fluorescence (mitochondria) with the endomembrane system (marked by mCherry fluorescence) were shown. Colocalization analysis was shown in C. Scale bars represent 10 μm.

**Table S1. Amino acid sequences of signal peptides for tested subcellular localizations used in this study.** The “xxxxxx” represents the extein residues after splicing.

| NUMBER | SUBCELLULAR LOCALIZATION SIGNALS | AMINO ACID SEQUENCES AFTER SPLICING |
|---|---|---|
| 1 | Mitochondrial Targeting Sequence (MTS) | MKFKAKFLTAWNNVKYGWVVKSRTSFSKIxxxxxxMKFKAKFLTAWNNVKYGWVVKSRTSFSKI |
| 2 | Nuclear Localization Signal (NLS) | KRPAATKKxxxxxxGQAKKKK |
| 3 | Plasma Membrane Localization Signal (PMS) | Hras: GCMSCKCVLS<br>Rac1: PVKKRKRKCLLL |
| 4 | CAAX | Hras: CVLS<br>Rac1: CLLL |

| | | |
|---|---|---|
| 5 | hGeminin Degron | MNPSMKQKQEEIKENIKNSSVPRRTLKMIQPSASGSLVGRENELSAGLSKRKHRNxxxxxxDHLTSTTSSPGVIVPESSENKNLGGVTQESFDLMIKENPSSQYWKEVAEKRRKAL |
| 6 | Skp1 Degron | PSIKLQSSDGEIFEVDVEIAKQSVTIKTMLEDLGMDDEGDDDPVPLPNVNAAILKKVIQWCTHHKDDPPPPEDDENKEKRTDDIPVWDQEFLKVDQGTLFELILAANYLDI |
| 7 | Tomm20 Mitochondrial target signal | MVGRNSAIAAGVCGALFIGYCIYFDRKRRSDPNFKNRLRERRKKQKLAKERAGLSKLPDLKDAEAVQKFFLEEIQLGEELLAQGEYEKGVDHLTNAIAVCGQPQQLLQVLQQTLPPPVFQMLLTKLPTISQRIVSAQSLAEDDVE |

**Table S2. Plasmids used in this study.**

| NUMBER | PLASMIDS | ANTIBIOTIC RESISTANCE | SOURCE |
|---|---|---|---|
| 1 | CSII-EF-mCherry-hGeminin_N-gp41-1_N | Amp+ | This study |
| 2 | CSII-EF-gp41-1_C-hGeminin_C | Amp+ | This study |
| 3 | CSII-EF-mCherry-hGeminin_N-NrdJ-1_N | Amp+ | This study |
| 4 | CSII-EF-NrdJ-1_C-hGeminin_C | Amp+ | This study |
| 5 | CSII-EF-mCherry-hGeminin_N-SspGyrB_N | Amp+ | This study |
| 6 | CSII-EF-SspGyrB_C-hGeminin_C | Amp+ | This study |
| 7 | CSII-EF-mCherry-NLS_N-gp41-1_N | Amp+ | This study |
| 8 | CSII-EF-gp41-1_C-NLS_C | Amp+ | This study |
| 9 | CSII-EF-mCherry-NLS_N-NrdJ-1_N | Amp+ | This study |
| 10 | CSII-EF-NrdJ-1_C-NLS_C | Amp+ | This study |
| 11 | CSII-EF-mCherry-NLS_N-SspGyrB_N | Amp+ | This study |
| 12 | CSII-EF-SspGyrB_C-NLS_C | Amp+ | This study |
| 13 | pcDNA3.1-AGGSAK-MTS-sfGFP | Amp+ | This study |
| 14 | pcDNA3.1-NPCSEI-MTS-sfGFP | Amp+ | This study |

| | | | |
|---|---|---|---|
| 15 | pcDNA3.1-SGYSSS-MTS-sfGFP | Amp+ | This study |
| 16 | pcDNA3.1-gp41-1_N | Amp+ | This study |
| 17 | pcDNA3.1-gp41-1_C-MTS-sfGFP | Amp+ | This study |
| 18 | pcDNA3.1-MTS_N-gp41_N | Amp+ | This study |
| 19 | pcDNA3.1-gp41_C-MTS_C-sfGFP | Amp+ | This study |
| 20 | pcDNA3.1-MTS_N-NrdJ-1_N | Amp+ | This study |
| 21 | pcDNA3.1-NrdJ-1_C-MTS_C-sfGFP | Amp+ | This study |
| 22 | pcDNA3.1-MTS_N-SspGyrB_N | Amp+ | This study |
| 23 | pcDNA3.1-SspGyrB_C-MTS_C-sfGFP | Amp+ | This study |
| 24 | pcDNA3.1-gp41-1_C-flag-mCherry | Amp+ | This study |
| 25 | pcDNA3.1-sfGFP-HNRNPA1a-gp41-1_N | Amp+ | This study |
| 26 | pcDNA3.1-sfGFP-HNRNPA1a-CRY2clust-gp41-1_ N' | Amp+ | This study |
| 27 | pcDNA3.1-CRY2clust-gp41-1_C'-flag-mCherry | Amp+ | This study |
| 28 | pcDNA3.1-sfGFP-FXR1a-cry2clust-gp41-1_N' | Amp+ | This study |
| 29 | pcDNA3.1-CRY2clust-flag-mCherry | Amp+ | This study |
| 30 | pcDNA3.1-gp41-1_C'-flag-mCherry | Amp+ | This study |
| 31 | pcDNA3.1-sfGFP-FXR1a | Amp+ | This study |
| 32 | pcDNA3.1-Skp1-CRY2-gp41-1_C'-Flag-mCherry | Amp+ | This study |
| 33 | pcDNA3.1-Skp1-gp41_1N'-CRY2 | Amp+ | This study |
| 34 | pcDNA3.1-sfGFP-gp41_1N'-CRY2 | Amp+ | This study |
| 35 | pcDNA3.1-CRY2-gp41-1_C'-Flag-mCherry-CVLS | Amp+ | This study |
| 36 | pcDNA3.1-CRY2-gp41-1_C'-Flag-mCherry-GCMS CKCVLS | Amp+ | This study |
| 37 | pcDNA3.1-CRY2-gp41-1_C'-Flag-mCherry-CLLL | Amp+ | This study |
| 38 | pcDNA3.1-CRY2-gp41-1_C'-Flag-mCherry-PVKK RKRKCLLL | Amp+ | This study |
| 39 | pcDNA3.1-TOM20-sfGFP-cry2-gp41-1N' | Amp+ | This study |
